# WNKs regulate mouse behavior and alter central nervous system glucose uptake and insulin signaling

**DOI:** 10.1101/2024.06.09.598125

**Authors:** Ankita B. Jaykumar, Derk Binns, Clinton A. Taylor, Anthony Anselmo, Sachith Gallolu Kankanamalage, Shari G. Birnbaum, Kimberly M. Huber, Melanie H. Cobb

**Author notes:** Co-corresponding author, 6001 Forest Park Rd., Dallas, Texas 75390-9041. Department of Pharmaceutical Sciences, College of Pharmacy, North Texas Eye Research Institute, UNT Health, Fort Worth, Texas- 76107, USA. Dana-Farber Cancer Institute, Center for Cancer Genomics, Boston, MA-02115, USA.

## Abstract

Certain areas of the brain involved in episodic memory and behavior, such as the hippocampus, express high levels of insulin receptors and glucose transporter-4 (GLUT4) and are responsive to insulin. Insulin and neuronal glucose metabolism improve cognitive functions and regulate mood in humans. Insulin-dependent GLUT4 trafficking has been extensively studied in muscle and adipose tissue, but little work has demonstrated either how it is controlled in insulin-responsive brain regions or its mechanistic connection to cognitive functions. In this study, we demonstrate that inhibition of WNK (With-No-lysine (K)) kinases improves learning and memory in mice. Neuronal inhibition of WNK enhances *in vivo* hippocampal glucose uptake. Inhibition of WNK enhances insulin signaling output and insulin-dependent GLUT4 trafficking to the plasma membrane in mice primary cortical neurons and hippocampal slices. Therefore, we propose that the extent of neuronal WNK kinase activity has an important influence on learning, memory and anxiety-related behaviors, in part, by modulation of neuronal insulin signaling.

## Background

Traditionally, the brain was considered an insulin-insensitive organ, based on whole-brain glucose uptake studies [1,2]. In contrast, more recent work has identified discrete areas of the brain, notably the hippocampus, cortex and the amygdala, that express high levels of insulin receptors and the insulin-sensitive glucose transporter glucose transporter-4 (GLUT4) [3–8]. In muscle and adipose tissue, insulin activates phosphatidylinositol 3-kinase (PI3K) and AKT (aka protein kinase B) signaling to cause the hallmark action of insulin in these tissues to increase GLUT4 translocation to the plasma membrane to facilitate insulin-dependent glucose uptake [9–10]. In a similar way, increases in neuronal insulin concentration also promote GLUT4 membrane translocation *via* PI3K/AKT signaling in neuronal cells [11–20]. Neuronal insulin signaling improves learning, memory and mood in humans and in animal models [18–29]. Glucose utilization increases in the hippocampus during memory tasks; performance on these challenging tasks is critically determined by an adequate supply of glucose to this brain region [30–42]. Recent studies show that hippocampal GLUT4 translocates to the plasma membrane during memory training. Further, acute intrahippocampal inhibition of GLUT4-mediated glucose transport impairs memory acquisition [18–19]. In addition, disrupted glucose metabolism in the hippocampus and other regions is associated with anxiety disorders [27–28, 37–38, 43–72]. These findings lead to the supposition that memory is enhanced, at least in part, *via* local neuronal insulin-dependent upregulation of glucose uptake to increase glucose available for metabolism.

The four WNKs (With-No-lysine (K) 1-4) are serine/threonine protein kinases best known for regulation of ion transport. One or more family members are responsible for rare hereditary diseases including forms of hypertension and sensory neuropathy [73–75]. The best characterized WNK substrates are the closely related kinases with overlapping functions that belong to the STE branch of the kinome tree: OSR1 (*OXSR1*, Oxidative stress response kinase 1) and SPAK (*STK39*, STE20/SPS-1-related proline-alanine-rich kinase) [76]. WNKs bind, phosphorylate, and activate OSR1 and SPAK which in turn phosphorylate downstream targets including the cation-chloride co-transporters NKCC1/2 (*SLC12A1/2*, Na^+^/K^+^/2Cl^-^ co-transporter), NCC (*SLC12A3*, Na^+^/2Cl^-^ co-transporter) and KCC2 (*SLC12A5*, K^+^/2Cl^-^ co-transporter) [76]. Genome-wide association studies have identified multiple SNPs in WNKs and several of these downstream effectors that are associated with impairment in learning, intellectual abilities, neuropsychiatric and age-related neurodegenerative diseases such as Alzheimer’s Disease (AD) etc. [77–94]. Abnormal activities of WNKs and their downstream effectors are also associated with these traits [95–109].

WNKs also regulate trafficking of many membrane transporters and ion channels [73, 110–119]. In addition to these well-characterized roles of WNKs, several studies have also suggested connected actions between WNK1 and AKT in cell lines and in kidneys including in hyperinsulinemic db/db mice [120–130]. PI3K/AKT signaling has been implicated in the regulation of memory and anxiety in mice [37,38]. These and other findings led us to hypothesize a critical function of the WNK pathway in memory and anxiety *via* regulating multiple steps in vesicle trafficking to modulate insulin-responsive GLUT4 translocation in neurons.

## Results

### Inhibition of hippocampal WNK in mice enhances learning and memory

WNK and its downstream signaling mediators, OSR1 and SPAK (**Figure 1A**), have been associated with learning and memory in animal models [77–90, 95–109]. mRNAs encoding WNKs 1-4 are expressed in areas of the brain such as the hippocampus based on mouse in situ hybridization data deposited in Allen Brain Atlas (**Figure S1A**). To examine the potential of WNKs on mouse learning and memory performance, we used the orally bioavailable pan-WNK inhibitor WNK463-this compound is selective for WNKs over other kinases based on a library of >400 protein kinases [131]. First, we evaluated the effectiveness of oral administration of WNK463 to inhibit WNKs in the mouse hippocampus. Phosphorylation of the WNK substrate OSR1 was a readout of WNK activity. Using this method, WNK463 modestly decreased pOSR1 (**Figure 1B, Figure S2A**) but had no adverse effect on mouse body weight (**Figure S3A**). We then tested the effect of WNK inhibition on memory performance by the Novel Object Recognition test. The Novel Object Recognition test takes advantage of the natural proclivity of mice to explore novelty; if a mouse recognizes a familiar object, it will spend more time exploring a novel object (**Figure 1C**). Therefore, longer time spent exploring a novel object compared to a familiar object (ie: a higher discrimination index [see methods for details]) is suggestive of better learning and/or memory. We found that the discrimination index for mice treated with oral WNK463 was significantly higher than that for vehicle-treated mice (**Figure 1D**), while the total exploration time during the test was similar between both groups (**Figure 1E**). We further tested the effect of WNK inhibition on hippocampal-dependent memory using the Contextual Fear Conditioning Test (**Figure 1F**). In the context test, WNK463-treated mice froze significantly longer than vehicle-treated mice, indicative of enhanced contextual memory in WNK463-treated mice (**Figure 1G**). However, no significant difference in freezing time was observed between the groups in the cue test, which assesses function of the amygdala, a major processing center of the brain for emotions such as fear (**Figure 1H**). To determine whether the increased freezing behavior might be due to an adverse effect of the drug or a confounding factor such as lethargy, we performed a locomotor test in a home cage-like environment (**Figure 1I**). We found no differences in their locomotor activity, making adverse effects unlikely to account for the results of the freezing test (**Figure 1J**). Results of these tests are consistent with the interpretation that WNK inhibition enhances learning and memory (**Figure 1K**).

**Figure 1.**
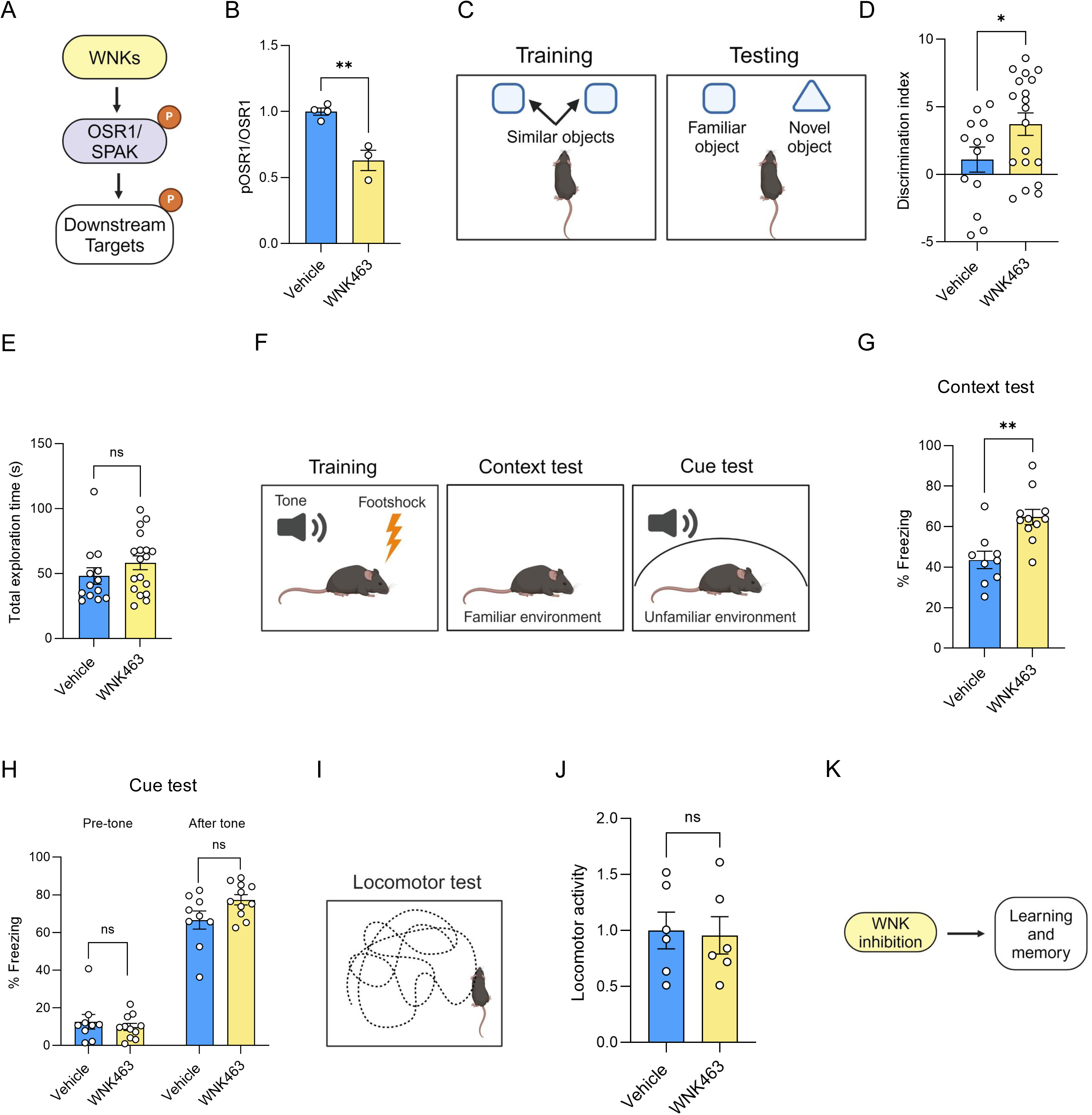
Inhibition of WNK in mice enhances learning and memory. **A)** Model shows WNK downstream signaling. **B)** Quantification shows pOSR1 (Ser325)/total OSR1 from hippocampi of mice treated with WNK463 (PO; 6 mg/kg) or vehicle; n=4. **C)** Diagram representing Novel Object Recognition test protocol. **D)** Quantification of discrimination index in mice administered with WNK463 (PO; 6mg/kg) (n=13) or vehicle (n=19) tested on the Novel Object Recognition protocol. **E)** Quantification of total exploration time during Novel Object Recognition test protocol in mice administered with WNK463 (PO; 6 mg/kg); n=13 or vehicle; n=19. **F)** Diagram representing Context-Cue Fear Conditioning test protocol. **G)** Quantification of % Freezing in Contextual Fear Conditioning test in mice administered WNK463 (PO; 6 mg/kg) (n=11) or vehicle: n=9. **H)** Quantification of % Freezing in Cued Fear Conditioning test in mice administered WNK463 (PO; 6mg/kg) (n=11) or vehicle: n=9. **I)** Diagram representing locomotor test protocol. **J)** Quantification of locomotory activity in mice administered WNK463 (PO; 6mg/kg) or vehicle; n=6. **K)** Model shows the effect of WNK inhibition on learning and memory in mice. Data are represented as Mean±SE; analyzed by unpaired two-tailed Student’s *t*-test or one-way ANOVA. ns: non-significant, *p<0.05, **p<0.005 and *** p<0.0005. Graphics created with BioRender.com.

### Inhibition of hippocampal WNK enhances anxiety-related behavior in mice

While the hippocampus is critical for cognitive processes such as episodic memory and spatial navigation, it is also implicated in the pathogenesis of anxiety disorders [18–28]. Several studies have confirmed a close correlation between anxiety and emotional memory [27–28, 37–38]. Enhanced fear-based contextual memory in WNK463-treated mice prompted us to test whether anxiety-like behaviors are altered in these mice. We found no difference in basal anxiety measured by the Open-Field test (**Figure 2A**, **2B**, **2C**) or the Elevated Plus Maze test (**Figure 2D**, **2E**, **2F**), between the vehicle and the WNK463-treated groups. However, stressing WNK463-treated mice with electric foot-shocks prior to the anxiety tests results in higher measures of anxiety. Mice treated with oral WNK463 spent less time in the center of the Open-Field test chamber relative to control mice, indicative of a higher level of anxiety (**Figure 2G**, **2H**). Similarly, these mice spent less time in the open arm of the Elevated Plus Maze compared to the control mice, also indicative of higher anxiety levels (**Figure 2I**, **2J**). Yet, the total distance moved by mice during both of these tests was not different between the groups (**Figure S4A, S4B, S4C, S4D**). This suggests that WNK463 only enhanced anxiety in mice pre-exposed to trauma such as electric foot-shocks.

**Figure 2.**
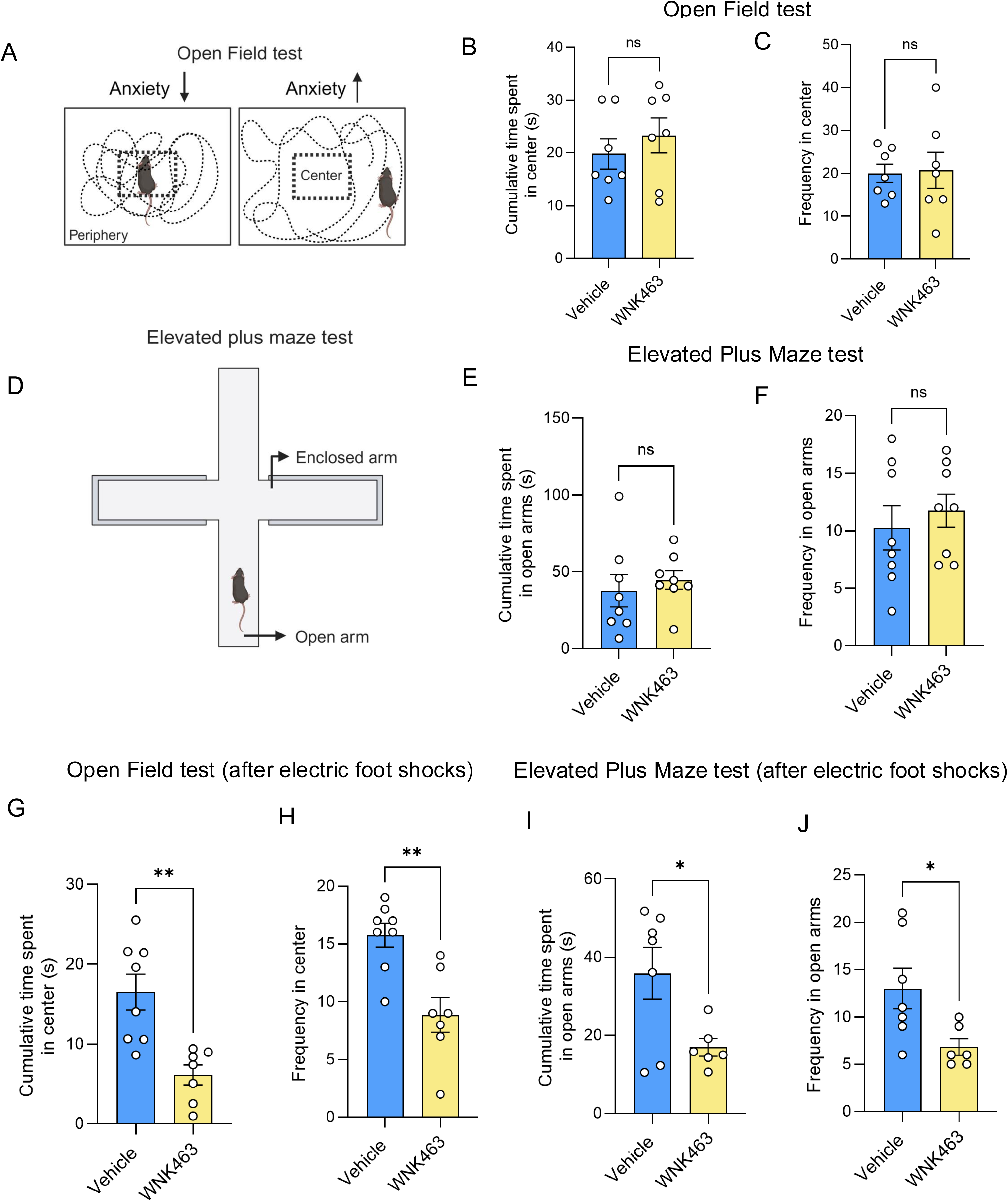
Inhibition of hippocampal WNK enhances anxiety-related behavior in mice. **A)** Diagram representing Open Field test protocol. **B)** Graph representing cumulative time spent in the center by mice treated with vehicle or WNK463 (PO; 6 mg/kg) in the Open Field test; n=7. **C)** Graph representing the number of entries (frequency) in the center by mice treated with vehicle or WNK463 (PO; 6 mg/kg) in the Open Field test; n=7. **D)** Diagram representing Elevated Plus Maze test protocol. **E)** Graph representing cumulative time spent in the open arms by mice treated with vehicle or WNK463 (PO; 6 mg/kg) in the Elevated Plus Maze test; n=8. **F)** Graph representing the number of entries (frequency) in the open arms by mice treated with vehicle or WNK463 (PO; 6 mg/kg) in the Elevated Plus Maze; n=8. **G)** Graph representing cumulative time spent in the center by mice treated with vehicle (n=8) or WNK463 (PO; 6 mg/kg) (n=7) in the Open Field test after mice received electric foot-shocks. **H)** Graph representing the number of entries (frequency) in the center by mice treated with vehicle (n=8) or WNK463 (PO; 6 mg/kg) (n=7) in the Open Field test after mice received the electric foot-shocks. **I)** Graph representing cumulative time spent in the open arms by mice treated with vehicle (n=7) or WNK463 (PO; 6 mg/kg) (n=6) in the Elevated Plus Maze test after mice received electric foot-shocks. **J)** Graph representing the number of entries (frequency) in the open arms by mice treated with vehicle (n=7) or WNK463 (PO; 6 mg/kg) (n=6) in the Elevated Plus Maze after mice received electric foot-shocks. Data are represented as Mean±SE; analyzed by unpaired two-tailed Student’s *t*-test or one-way ANOVA. *p<0.05, **p<0.005 and *** p<0.0005. ns: non-significant; p>0.05.

### Inhibition of WNK augments glucose uptake via GLUT4

Insulin action and glucose uptake into select regions of the brain are thought to regulate memory and anxiety-like behavior in mice [18–32, 37–38, 43–72]. We found oral administration of WNK463 in mice did not affect fasting serum insulin levels (**Figure S5A**). Because WNK inhibition enhanced memory performance and anxiety in mice, we tested whether inhibition of WNKs in insulin-sensitive areas of the brain enhanced 2-deoxyglucose uptake using an *in vivo* radioactive 2-deoxy glucose uptake assay in mice. Dissecting the hippocampi of mice treated with WNK463, we found 2-deoxyglucose uptake in this insulin-sensitive region was elevated compared to hippocampi from mice treated with vehicle (**Figure 3A**). These data together suggested that inhibition of hippocampal WNK enhanced *in vivo* hippocampal glucose uptake in mice. Next, we cultured hippocampal slices from mouse brain, and found that inhibition of WNK enhanced 2-deoxyglucose uptake in these hippocampal slice cultures (**Figure 3B**). We also isolated crude synaptosomes from mouse brain and found that inhibition of WNK enhanced 2-deoxyglucose uptake in these synaptosomes (**Figure 3C**). To examine the effect of insulin in cultured cells, SH-SY5Y cells were differentiated to neuronal-like cells *in vitro* [132]. Stimulation of these cells with insulin induced [^3^H]-2-deoxyglucose uptake that was further enhanced by simultaneous inhibition of WNKs. In addition, WNK463 alone enhanced 2-deoxyglucose uptake compared to DMSO control (**Figure 3D**).

**Figure 3.**
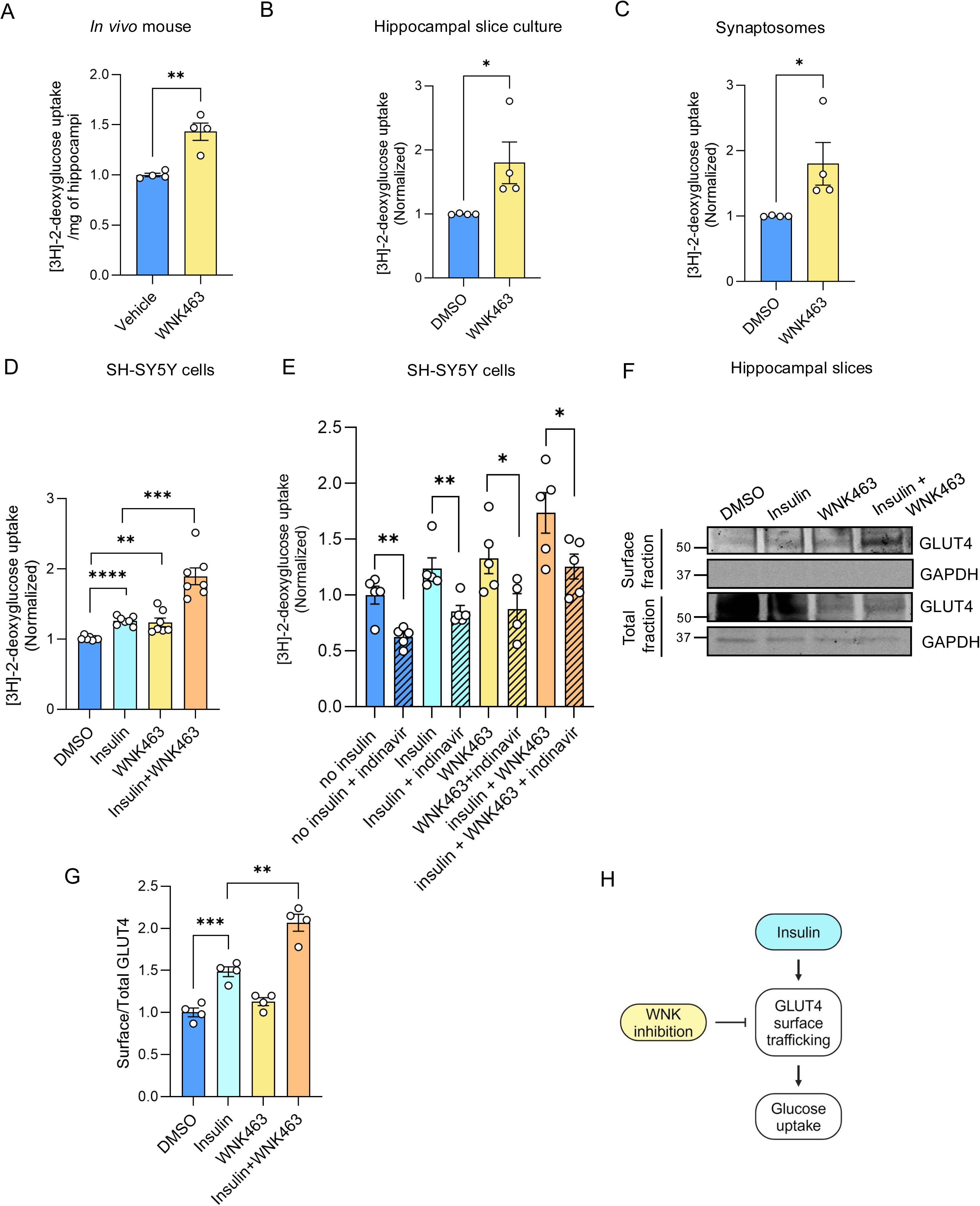
Inhibition of WNK augments glucose uptake via GLUT4. **A)** Graph representing quantification of *in vivo* radioactive 2-deoxyglucose uptake per mg hippocampal weight in mice treated with vehicle or WNK463 (PO; 6 mg/kg); n=4. **B)** Graph representing enhanced radioactive 2-deoxyglucose uptake in hippocampal slice culture from C57BL/6J mouse treated with WNK63 (1 µM); n=4. **C)** Graph representing enhanced radioactive 2-deoxyglucose uptake in crude synaptosome from C57BL/6J mouse whole brains treated with WNK63 (1 µM); n=3. **D)** Graph shows *in vitro* radioactive 2-deoxyglucose uptake in SH-SY5Y cells treated with WNK463 (1 µM), insulin (10 nM); n=7. **E)** Graph representing enhanced radioactive *in vitro* 2-deoxyglucose uptake in differentiated SH-SY5Y cells treated with insulin (10 nM) ± WNK463 (1 µM) ± indinavir (10 nM); n=5. **F)** Representative Western blot shows surface and total GLUT4 protein fraction from mice hippocampal slices treated with insulin (10 nM) and/or WNK463 (1 µM). **G)** Corresponding quantification of ‘G’ shows enhanced surface GLUT4 (measured as a fraction of total GLUT4) with WNK463 ± insulin treatment; n=4. **H)** Model shows downstream effects of insulin and the impact of WNKs on insulin signaling. Data are represented as Mean±SE; analyzed by unpaired two-tailed Student’s *t*-test or one-way ANOVA. *p<0.05, **p<0.005 and *** p<0.0005. Graphics created with BioRender.com.

Because insulin regulates GLUT4, we asked whether increased glucose uptake in WNK-inhibited neuronal cells depends on GLUT4 [70–72]. Differentiated SH-SY5Y cells were treated with indinavir, a GLUT4 inhibitor, to identify the contribution of GLUT4 to enhanced uptake caused by WNK463. Indinavir reduced the enhanced insulin-stimulated 2-deoxyglucose uptake observed with WNK inhibition, suggesting a role for GLUT4 in facilitating at least a fraction of glucose uptake responsive to WNK blockade (**Figure 3E**). To obtain further evidence for the involvement of GLUT4, we examined GLUT4 localization in mice hippocampal slices using surface biotinylation. WNK inhibition with WNK463 enhanced insulin-stimulated GLUT4 surface expression relative to insulin alone (**Figure 3F**, **3G**). We obtained similar results using differentiated SH-SY5Y cells for surface biotinylation in the presence of WNK463, insulin or both (**Figure S6A, S6B**). Together, these data suggest that inhibition of WNK in neuronal cells enhances insulin-dependent glucose uptake via enhanced GLUT4 surface targeting (**Figure 3H**).

### Inhibition of WNK enhances AKT signaling in the hippocampus and neuronal cell culture

Insulin induces activation of PI3K/AKT signaling to cause GLUT4 translocation to the plasma membrane to facilitate insulin-dependent glucose uptake [9–10] Given the involvement of insulin in regulating GLUT4 trafficking, we asked whether inhibition of WNK in neuronal cells affects insulin signaling to PI3K/AKT. Multiple groups including ours have previously reported inhibitory cross talk between WNK1 and AKT [120–130]. In homogenates of excised hippocampal tissue from mice treated with oral WNK463 compared to vehicle, we found enhanced phosphorylation of AKT (pAKT- at Ser473 which is an activating phosphorylation site on AKT) (**Figure 4A**, **4B**), indicative of enhanced insulin signaling. In differentiated SH-SY5Y neuroblastoma cells, we found that co-treatment with insulin and the WNK inhibitor enhanced pAKT compared to insulin alone (**Figure 4C**, **4D**). This is consistent with enhanced insulin signaling resulting from WNK inhibition in other cell lines we tested (HeLa- **Figure S7A, S7B,** HUVECs**- S7C, S7D,** SH-SY5Y**- S7E, S7F**). In dissociated mouse primary cortical neurons, we found that WNK inhibition enhanced pAKT compared to vehicle control (**Figure 4E**, **4F**), although in this case, WNK inhibition and insulin co-treatment failed to further enhance activation of AKT compared to insulin alone (**Figure 4G**).

**Figure 4.**
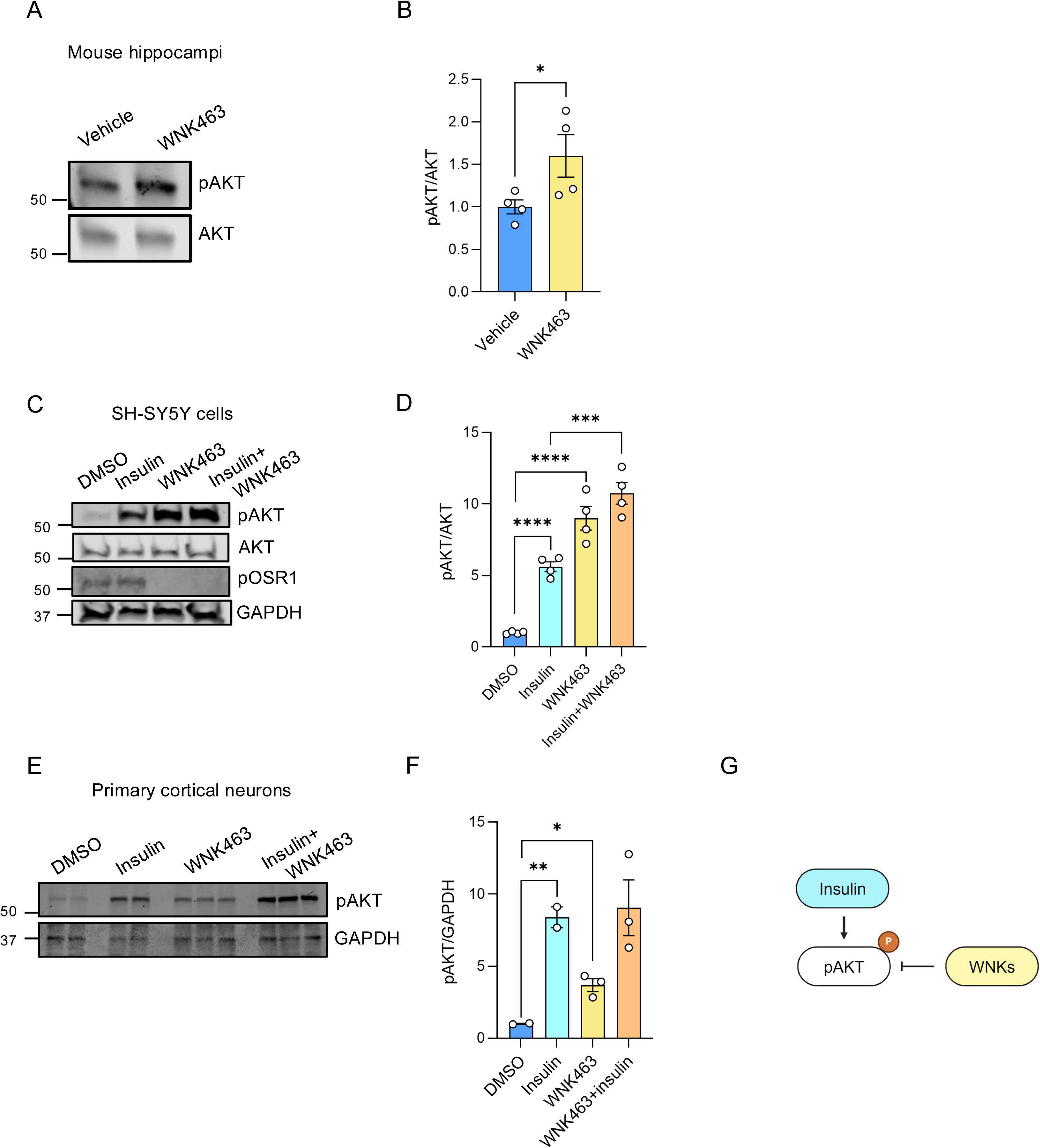
Inhibition of WNK enhances insulin signaling in the hippocampus and cell culture. **A)** Representative Western blots showing pAKT, AKT levels in excised hippocampal tissue from mice treated with vehicle or WNK463 (PO: 6 mg/kg) for 3 days; n=4**. B)** Quantification of ‘B’. **C)** Representative Western blot shows pAKT, AKT and pOSR1 in the presence of WNK463 (1 µM) ± insulin (10 nM) in SH-SY5Y cells. **D)** Quantification of pAKT/AKT in differentiated SH-SY5Y cells treated with insulin (10 nM) ± WNK463 (1 µM); n=4. **E)** Representative Western blot shows pAKT and GAPDH in presence of WNK463 (1 µM) ± insulin (10 nM) in mouse primary cortical neurons. **F)** Quantification of pAKT/GAPDH in primary cortical neurons treated with insulin (10 nM) ± WNK463 (1 µM); n=2 for DMSO and insulin and n=3 for WNK463±insulin. **G)** Model shows insulin signaling and the impact of WNK kinases on insulin signaling. Data are represented as Mean±SE; analyzed by unpaired two-tailed Student’s *t*-test or one-way ANOVA. *p<0.05, **p<0.005 and *** p<0.0005. Graphics created with BioRender.com.

### OSR1/SPAK interact with molecular mediators of GLUT4 trafficking

Upon insulin signaling, AKT phosphorylates the AKT-substrate of 160-kDa (AS160, *TBC1D4*) [116–118], a Rab GTPase-activating protein critical in liberating the static pool of GLUT4 from its storage vesicle site to promote its exocytosis [117–118]. Studies conducted by the Jordan lab suggested that WNK1 binds to AS160 and subsequently identified WNK1 phosphorylation sites in the protein [116–117]. OSR1 and SPAK are two-domain enzymes containing a kinase domain and a conserved carboxy-terminal (CCT) domain that houses a docking site for proteins containing short basic/hydrophobic motifs, subsequently confirmed through structural analysis [76]. Protein interaction studies showed that the motif, most typically R-F-x-V/I or R-x-F-x-V/I often forms the basis for substrate recognition [110,133]. Bioinformatic analysis of motif interactions with the CCT, returned AS160 as a likely OSR1/SPAK binding protein through its R-x-F-x-I motif [134]. Thus, we asked whether the WNK/OSR1/SPAK pathway regulates GLUT4 trafficking [119], in part by modulating AS160 function. First, we found that endogenous AS160 from mouse brain lysates co-immunoprecipitated with OSR1 (**Figure 5A**). To ask whether the interaction between AS160 and OSR1 is modulated by insulin, we immunoprecipitated endogenous AS160 from differentiated SH-SY5Y cells treated with or without insulin and quantified the amount of OSR1 that co-immunoprecipitated. We found that insulin treatment enhanced the interaction between AS160 and OSR1, suggesting that insulin influences the OSR1-dependent effect on AS160 localization. We also found that co-treatment with WNK463 reduced the interaction between AS160 and OSR1 (**Figure 5B**, **5C**). Insulin regulates AKT-dependent phosphorylation of AS160 on Ser588. We found enhanced phosphorylation of Ser588 on AS160 following WNK inhibition consistent with the increase in AKT activity caused by suppressing WNK activity (**Figure 5D**, **5E**).

**Figure 5.**
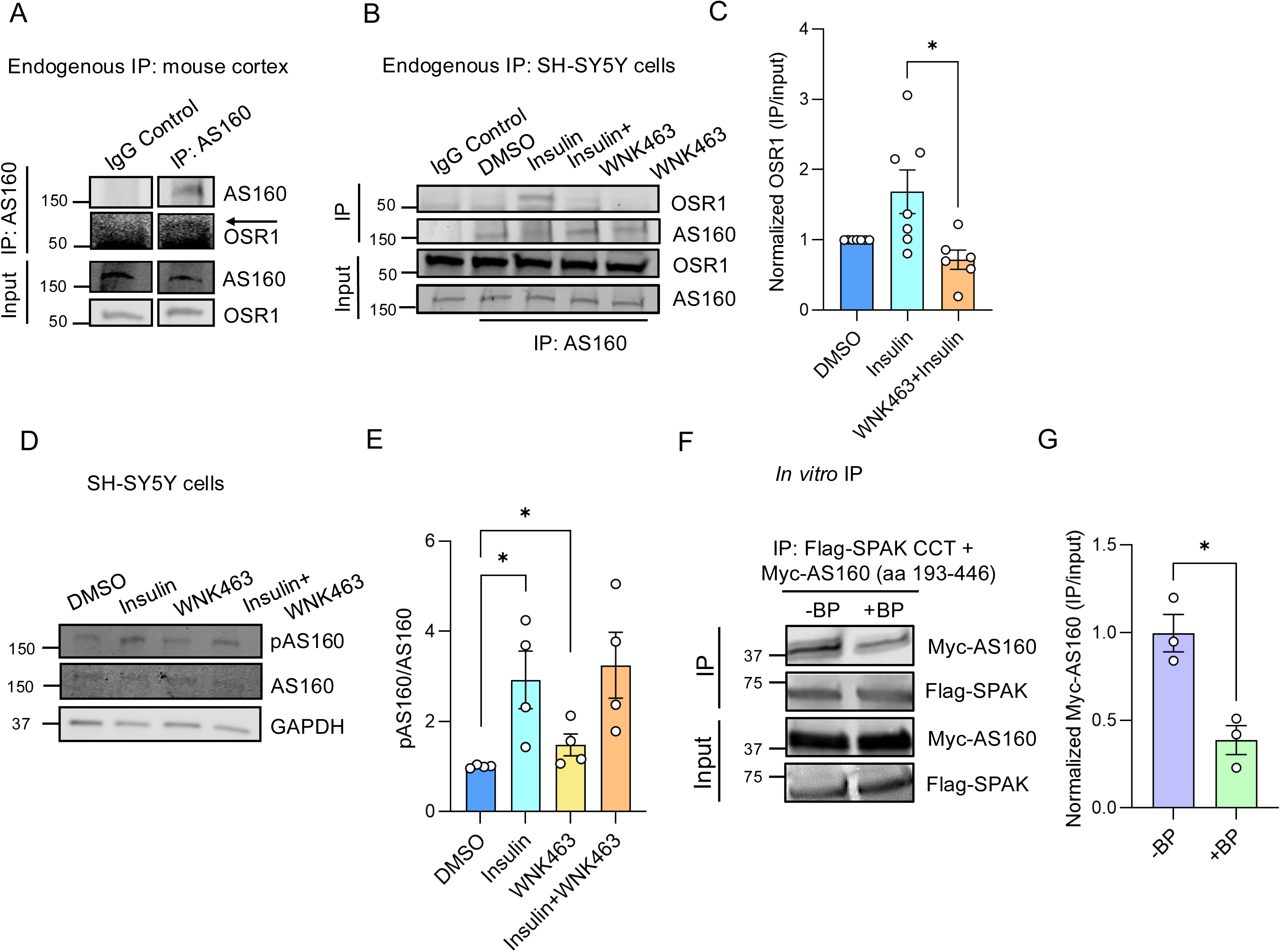
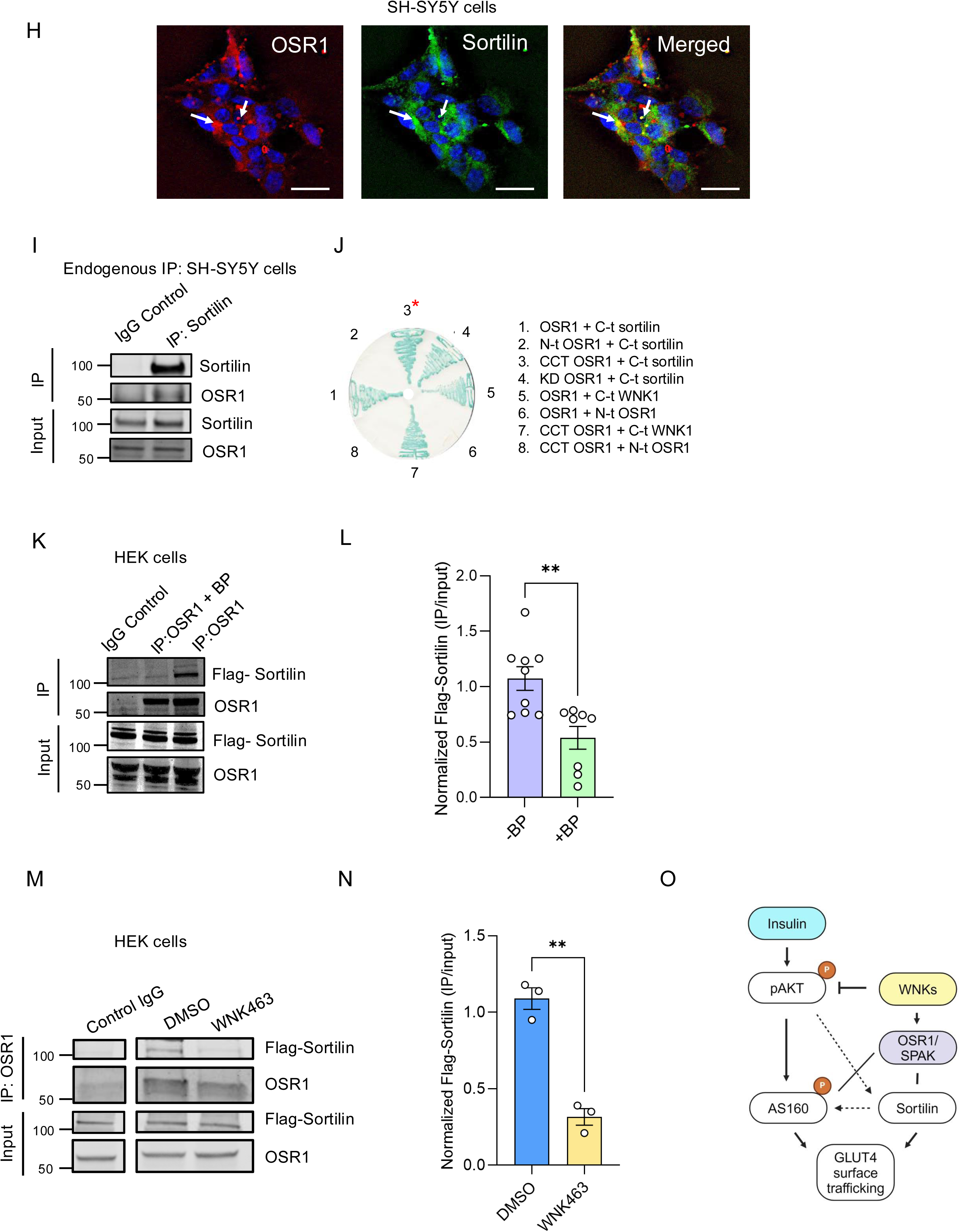
OSR1 interacts with molecular mediators involved in GLUT4 trafficking. **A)** Representative Western Blot shows co-immunoprecipitation of endogenous OSR1 with AS160 from C57BL/6J whole mouse brain lysates; n=3. **B)** Representative Western blot shows co-immunoprecipitation of AS160 with OSR1 upon treatment with WNK63 (1 µM) and/or insulin (10 nM) in differentiated SH-SY5Y cells. **C)** Corresponding quantification of ‘**B**’ shows decreased association of AS160 and OSR1 in cells treated with insulin (10 nM) + WNK463 (1 µM) compared to insulin alone; n=6. **D)** Representative Western blot shows pAS160, AS160 and GAPDH in differentiated SH-SY5Y cells treated with insulin (10 nM) ± WNK463 (1 µM). **E)** Corresponding quantification of ‘D’ shows increased pAS160 in cells treated with insulin (10 nM) or WNK463 (1 µM) compared to DMSO; n=4. **F)** R-x-F-x-V- containing blocking peptide (BP: WNK1 1253-1265; NH_3_^+^-SAGRRFIVSPVPE-COO^-^; 100 µM) decreases interaction of overexpressed OSR1/SPAK CCT bait protein fragment (aa 50-545) with myc-AS160 protein fragment (aa 193-446) *in vitro*; n=3. **G)** Corresponding quantification of ‘F’. **H)** Bright-field (left) or confocal images shows co-localization of endogenous OSR1 with sortilin in differentiated SH-SY5Y cells. OSR1 (red), sortilin (green), merged (yellow), scale bar=10 µm; n=3. **I)** Representative endogenous co-immunoprecipitation of OSR1 with sortilin in differentiated SH-SY5Y cells; n=5. **J)** Yeast two-hybrid assay shows binding of the conserved C-terminus (CCT) of OSR1 with the C-terminus (C-t) of sortilin (condition 3- denoted by *); binding of full-length and CCT of OSR1 to the C-terminus of WNK1 as positive controls (5,7). N-t: N-terminus. **K)** Representative Western blot shows co-immunoprecipitation of OSR1 and Flag-sortilin in HEK cells is diminished upon co-incubation with the blocking peptide (BP) SAGRRFIVSPVPE; n=3. **L)** Corresponding graphical representation for ‘K’; n=8. **M)** Representative Western Blot shows co-immunoprecipitation of OSR1 and Flag-sortilin in HEK cells is diminished upon WNK463 treatment; n=3. **N)** Corresponding quantification of **‘M’. O)** Model shows balanced regulation of sortilin and AS160 by the WNK/OSR1/SPAK pathway to regulate GLUT4 trafficking. Data are represented as Mean±SE; analyzed by unpaired two-tailed Student’s *t*-test or one-way ANOVA. *p<0.05, **p<0.005 and *** p<0.0005. Graphics created with BioRender.com.

Using an *in vitro* pull-down assay with purified SPAK isolated from Human Embryonic Kidney (HEK293T) cells, 3xFlag-SPAK (residues 50-545), co-immunoprecipitated with a fragment of AS160 (residues 193-437) containing the R-x-F-x-V motif. Moreover, this interaction was diminished by co-incubation with a blocking peptide derived from WNK1 (SAGRRFIVSPVPE; residues 1253-1265) that competes for docking interactions on the OSR1/SPAK CCT domain (**Figure 5F**, **5G**). This suggests that AS160 associates with SPAK during insulin signaling. Thus, the WNK/OSR1/SPAK pathway likely communicates with this step of the GLUT4 translocation process via direct interaction of AS160 with OSR1/SPAK and possibly through enhancing AKT activity to phosphorylate AS160. These data reveal the potential of WNK/OSR1/SPAK to influence insulin-sensitive GLUT4 trafficking at this essential step of the process.

Optimal GLUT4 trafficking to the membrane in response to insulin treatment is contingent on its efficient sequestration in specialized GLUT4 storage vesicles, a phenomenon common to both neurons and peripheral insulin-sensitive tissues [9–11]. The protein sortilin (*SORT1*, also known as Neurotensin receptor-3) is essential for the formation of and retrieval of GLUT4 to these static storage vesicles in the trans-Golgi network (TGN) which allows efficient GLUT4 translocation to the plasma membrane upon insulin treatment [11,135–139]. Sortilin-mediated GLUT4 retrograde trafficking and sequestration is critically dependent on a highly conserved motif with the consensus R-F_787_-L-V in sortilin [70–71,140]. However, the underlying molecular mediators that regulate this crucial process are unknown. Our motif bioinformatics also predicted that the R-F-x-V motif in the C-terminus of sortilin binds to the OSR1/SPAK CCT domain and raised the possibility that this plays a critical role in regulating retrograde trafficking of GLUT4 to the TGN [110,133]. Therefore, we asked whether WNK/OSR1/SPAK interacts with sortilin. We found that OSR1 and sortilin colocalize in differentiated SH-SY5Y cells (**Figure 5H**). We also found that endogenous OSR1 and sortilin co-immunoprecipitate from differentiated SH-SY5Y cells (**Figure 5I**). Previously, a yeast-two-hybrid screen using the OSR1 CCT domain as bait had returned sortilin as a potential binding partner (**Figure 5J**). To determine whether or not the interaction between the CCT domain of OSR1 and sortilin is facilitated by its R-F-x-V motif, we overexpressed sortilin in HEK293 cells and immunoprecipitated it. We found that OSR1 was also recovered in the sortilin precipitate. Co-immunoprecipitation of OSR1 with sortilin is prevented by co-incubation with the blocking peptide that competes with the R-F-x-V docking motif for binding to the OSR1 CCT (**Figure 5K**, **5L**). As a further confirmation, we tested the binding affinities of wild-type sortilin and sortilin R-F-x-V motif mutant peptides with bacterially expressed purified His_6_-SPAK CCT domain (residues 433-527). We found that only the WT sortilin peptide bound strongly to the CCT domain, while substitution of the motif R with K or A led to a drastic reduction in binding affinity. This implies that the interaction between sortilin and OSR1/SPAK CCT domain is facilitated via the key sortilin R-F-x-V motif (**Figure S8A**). We find that WNK463 blocks co-immunoprecipitation of Flag-tagged Sortilin with endogenous OSR1 in HEK293T cells (**Figure 5M**, **5N**). These findings lead to the idea that WNK has the potential to impact this essential step in GLUT4 trafficking through binding of its effector kinases OSR1/SPAK to sortilin. Together, our results further suggest that WNK/OSR1/SPAK likely influences insulin-sensitive GLUT4 trafficking by balancing GLUT4 sequestration in the TGN via regulation of sortilin with GLUT4 release from these vesicles upon insulin stimulation via regulation of AS160 (**Figure 5O**).

## Discussion

Insulin sensitivity is highly dependent upon confinement of GLUT4 in specialized GLUT4 storage vesicles until insulin signals a release from these vesicles to promote GLUT4 membrane translocation [9–11]. In insulin-resistant states, derangements in the formation and retention of GLUT4 in these static vesicles impairs insulin-responsive GLUT4 translocation [72]. In peripheral tissues, insulin resistance is associated, in part, with reduced glucose transport due to a loss-of-function in insulin-regulated GLUT4 [63]. Underlying mechanisms in the brain, however, remain unclear. Here we evaluated two proteins, sortilin and AS160, important for GLUT4 trafficking (Figure 3H, 5O). Sortilin which is widely expressed in neurons is suggested to be a critical mediator of this process and hence an important determinant of neuronal insulin sensitivity [11,135–139,141]. AS160 is phosphorylated by AKT upon insulin stimulation and causes the release of GLUT4 trapped in the static GLUT4 storage vesicles leading to exocytosis and surface delivery of GLUT4 to promote glucose uptake into cells [118]. In this study, we demonstrate actions of the WNK pathway on these two GLUT4 trafficking mediators sortilin and AS160. Our study shows that WNKs could affect both capture of GLUT4 in storage vesicles via sortilin as well as the release of trapped GLUT4 from the storage vesicles to facilitate glucose uptake into the cells via AS160. This work reveals at least two sites of action of the WNK pathway on a hallmark action of insulin- translocation of GLUT4 to the plasma membrane.

We suggest that the downstream effector kinases OSR1/SPAK have positive and perhaps required actions in this pathway. OSR1 appears to be needed for proper sequestration of GLUT4 in, and possibly release of GLUT4 from the storage vesicles. As a result, overall, the WNK pathway appears to be a positive regulator of GLUT4 trafficking. This is consistent with results of two other studies from different laboratories. The first study in skeletal muscle shows that loss of WNK1 (*rather than inhibition of activity*) drastically reduced GLUT4 localization to the plasma membrane in response to insulin, suggesting that WNK is essential for this trafficking event [119]. The second study shows that overexpression of the C-terminal tail of sortilin containing the R-F-x-V motif had an inhibitory effect on GLUT4 trafficking [142], suggesting that blocking the action of OSR1/SPAK prevents proper GLUT4 sequestration in storage vesicles. We anticipate that in the absence of a stimulus or a WNK inhibitor, low level WNK activity is sufficient to support OSR1/SPAK function to permit some of the trafficking functions of these proteins, including aiding the basal function of sortilin and GLUT4 assembly in storage vesicles, which may also involve AKT [142]. Then, activation of AKT by insulin will release GLUT4 to the membrane, independent of a substantial change in WNK activity. If this were not the case, only in cells in which WNKs were highly active, could OSR1 function as a component of the sequestration step or the transfer mechanism mediated by AS160 as a result of insulin-stimulated AS160-OSR1 binding. Chemical WNK inhibition will enhance AKT to increase its transporter localization to the plasma membrane. However, under normal conditions, in the absence of an inhibitor, basal/low level WNK activity will be adequate to permit the sortilin get-ready steps in vesicle formation and insulin/AKT will do the rest. We are working to establish the validity of this proposed explanation.

In addition to results presented here with WNK inhibition, several lines of evidence suggest cross-talk among AKT and WNK signaling pathways [120–130]. One early study suggested that insulin induces phosphorylation of the WNK downstream target, NCC (Na^+^/2Cl^-^ cotransporter) in mouse kidney cells [123]. WNK1 is a substrate of AKT which phosphorylates it on T60 [124–125] but phosphorylation on this residue has not been shown to affect WNK catalytic activity, although several studies have used WNK1 pT60 as a proxy for its function [119, 143]. In mouse embryonic fibroblasts it was reported that phosphorylation of WNK1 T60 by AKT promotes its degradation via the ubiquitin-proteasome pathway [129]. Phosphorylation of WNK1 on T60 contributes to the activation of the Serum and Glucocorticoid Regulated kinase 1 (SGK1) in a PI3K-dependent manner [127–128]. WNK1/SGK1 signaling is chronically activated in the AKT3 knockout mice model, contributing to high fat diet-induced metabolic syndrome, and this was reversed upon SGK1 inhibition [129–130]. The PI3K/AKT pathway activates the WNK-OSR1/SPAK/NCC cascade in hyperinsulinemic db/db mouse, model of insulin resistance [122]. In contrast, another study reported diminished insulin/AKT/WNK1 functions in diabetic skeletal muscle [119]. Although mechanistic clarity is lacking, together these findings suggest a critical role of aberrant WNK1 signaling in the development of metabolic syndrome. In this study, we show that WNK inhibition using the inhibitor WNK463 enhances insulin/AKT signaling, suggesting a reciprocal cross-talk between WNK pathway and insulin/AKT signaling. Interestingly, db/db mice exhibit progressive age-dependent decrease in AKT activity, surface GLUT4 as well as dysregulated WNK signaling in the hippocampus and this impairment occurs prior to the overt development of cognitive decline in this mouse model [66–69,144–154]. Interventions that correct dysregulated AKT downstream function ameliorated cognitive impairment in db/db mice [151,155,156]. Thus, we propose that aberrant WNK signaling can impact cognitive decline due to insulin/AKT metabolic dysregulation.

Sortilin is a member of the family of vacuolar protein sorting 10 protein (VPS10P) domain receptors. In addition to its glucose transport function, it has multiple actions in neurons [157–159]. Genetic and functional studies implicate sortilin and sortilin-related receptor 1 in aging, age-related cognitive decline, and AD [157–164]. Mechanistically, sortilin is involved in trafficking of BDNF, implicated in memory processes [158]. It also mediates endocytic uptake of Apolipoprotein E (APOE)-bound amyloid-β-containing lipoproteins in neurons [159]. The APOE-(4) isoform of APOE bears significance as the major genetic risk factor for late-onset AD [165–168]. Functionally, the APOE4 isoform perturbs sortilin trafficking [93] and impairs neuronal insulin signaling by disrupting insulin receptor trafficking [169]. Humans carrying the APOE4 isoform exhibit compromised glucose metabolism and downregulation of GLUT4 expression in the hippocampus, probably via affecting sortilin trafficking [170]. Our study suggests that inhibition of WNK kinases promotes glucose uptake in mice hippocampi and regulate hippocampus-dependent memory. Mechanistically, this could be, in part, due to regulation of sortilin (Figure 3H, 5O).

Many studies have linked abnormal WNK activity to an array of neurological diseases. Several SNPs, mutations and aberrant activities in WNK pathway components have been associated with learning disabilities, neuropsychiatric and neurodegenerative diseases [77–109,91–94]. WNK1 is highly expressed in the CA1-CA3 regions of the dorsal hippocampus, implicated in contextual memory [171]. WNK1 is differentially expressed in the hippocampus from schizophrenic and AD patients [101–102]. Several pathogenic variants in the *WNK3* gene cause intellectual disability [94]. Several mutations in the *WNK3* gene were identified in schizophrenic patients and expression of WNK3 protein was also elevated in these patients [99,103,172]. WNK3 regulates the splicing factor Fox-1 implicated in anxiety disorder and schizophrenia [107,173]. Activities of some well-characterized downstream effectors of the WNK pathway, e.g., OSR1, SPAK, NKCC1 and KCC2, are altered in the hippocampus in multiple models of neuropsychiatric diseases and neurodegenerative diseases [95–106]. The control of ion homeostasis has been assumed to be the mechanism underlying the above-mentioned neuro-pathophysiologies. Our study adds insights on the possible underlying causes of WNK pathways in regulating neuronal functions.

Aging is a major risk factor for the development of diabetes and severe insulin resistance [174,175]. Neuronal insulin resistance marked by striking reductions in insulin signaling and disrupted hippocampal glucose uptake are early indicators of cognitive decline in age-related diseases such as Alzheimer’s Disease (AD) [43–62]. Interestingly, the neurocognitive impairment commonly known as HIV-associated neurocognitive disorder is prevalent in HIV patients administered anti-retroviral GLUT4-impairing protease inhibitors (indinavir, nelfinavir) compared to patients treated with GLUT4-sparing protease inhibitors (such as Atazanavir) [64–65]. Animal models of insulin resistance exhibit diminished hippocampal plasma membrane GLUT4 [63–67] and correction of insulin resistance rescued the impairment in hippocampal-dependent memory and synaptic plasticity [68–69]. Therefore, several lines of research converge to suggest that aberrant insulin-responsive GLUT4 translocation causes hippocampal cognitive dysfunction, an overlap between metabolic and cognitive disorders. Our study suggests that WNK kinases are critical mediators of insulin sensitivity in neurons which affects learning and memory. Aberrant regulation of these kinases could disrupt normal neuronal behavior. Therefore, an important direction for future studies will be to determine how WNK regulates cognitive behavior in neuronal insulin resistant states.

## Methods

### Cell lines

The human neuroblastoma cell line (SH-SY5Y: ATCC, CRL-2266) was grown in low glucose DMEM medium (11885084, Thermo Fisher Scientific) with 10% fetal bovine serum (Sigma-Aldrich, F0926), 1% L-glutamine, 1% penicillin and streptomycin. SH-SY5Y cells were differentiated in Neurobasal medium (21-103-049, Fisher Scientific) with Glutamax^TM^ (35-050-061, Fisher Scientific), B27 supplement (17504044, Thermo Scientific), 1% penicillin and streptomycin along with 10 µM retinoic acid (R2625, Millipore Sigma) [132]. Alternatively, for glucose uptake assays, these cells were differentiated in low glucose DMEM (Thermo Fisher Scientific, 11885084) with 10 µM retinoic acid. Human embryonic kidney cells (HEK293; CRL-1573), Human Umbilical Vein Endothelial Cells (HUVECs) and HeLa were purchased from ATCC and were grown in DMEM medium with 10% FBS and 1% penicillin-streptomycin. All the cells were maintained at 37°C and 5% CO_2_.

### Dissociated mouse primary cortical neurons and glial cell culture

Dissociated cortical cultures were prepared from P0 mice using modified, previously published protocols [176]. Briefly, dissected cortex from P0 mice were trypsinized for 10min then dissociated by trituration. After centrifugation, neurons were plated in Neurobasal A medium (Invitrogen) containing B27 (2%; Invitrogen), 0.5 mM Glutamax, 1% Pen-Strep, and 5% fetal bovine serum (FBS) at a density of 8×10^5^ neurons per well of a 12-well dish each coated with 1 mg/ml poly-L-lysine overnight. Day 2, the medium was changed to Neurobasal A medium (Invitrogen), B27 (2%; Invitrogen), 0.5 mM Glutamax, without FBS. Day 3, cytosine arabinoside (1 µM) was added. Day 5, plates were washed 1 × with Neurobasal A with B-27 (Life Technologies) and 0.5 mM Glutamax and replaced with glial conditioned Neurobasal A medium containing B27, glutamine and cytosine arabinoside (1 μM). Cultures were fed every 5 days by replacing 50% of the medium with glial conditioned medium. At day 21, neurons were treated as described.

Glial cultures were prepared from the neocortex of P0-P2 mouse pups and maintained in Neurobasal A containing 10% FBS, 2% B27 supplement and 50 μg/ml penicillin, 50 U/ml streptomycin, Sigma). Medium was replaced twice a week for 2-3 weeks. Cells were conditioned in Neurobasal medium lacking B27 and FBS for 48hr, collected and stored at 4°C for no more than one week prior to addition to neuronal cultures.

### Hippocampal slices

Hippocampal slices (400 µm) were prepared from 30-45-day-old C57BL/6J mice as in [177]. Mice were anesthetized with xylaxine (4mg/ml)/ketamine (30 mg/ml). Mice were subjected to transcardial perfusion with ice-cold dissection buffer containing: 121 mM choline chloride, 2.5 mM KCl, 1.25 mM NaH_2_PO_4_, 5 mM dextrose, 30 mM NaHCO_3_, 3 mM ascorbic acid and adjusted to 290 mOsm. Mice were decapitated, and the cerebrum was dissected from isolated brains and then sliced using a vibratome (VT 1000S; Leica, Nussloch, Germany) in the same ice-cold dissection buffer. The slices were transferred into a reservoir chamber filled with artificial cerebrospinal fluid (aCSF) containing 119 mM NaCl, 2.5 mM KCl, 31 mM NaHCO_3_, 5 mM D-glucose, 1 mM NaH_2_PO_4_ [177]. Final concentrations of 1 mM MgCl_2_ and 2 mM CaCl_2_ were added just before use. Slices were allowed to recover for 2-3hr at 30°C. Both the aCSF and the dissection buffer were equilibrated with 95% O_2_ and 5% CO_2_.

### Plasmid constructs

Sortilin full-length (NM_001205228) construct was cloned into a C-terminally tagged 3xFlag CMV14 vector (Sigma Aldrich) with restriction enzymes XbaI. Clones were screened by XbaI digests. Positive clones were verified by Sanger sequencing.

### Yeast Two-Hybrid Analysis

A Jurkat T cell cDNA library (from Mike White, formerly Department of Cell Biology, University of Texas Southwestern Medical Center) was screened as described in [178]. Protein–protein interactions were tested by streaking co-transformants on medium lacking Leu, Trp, and His in addition to β-galactosidase assays.

### Co-immunoprecipitation

Cells were lysed in lysis buffer (50mM HEPES, 150 mM NaCl, 5mM EDTA, 2% Triton X-100, 0.1% SDS) with 1:1000 protease inhibitor cocktail (stock containing: 583.2 µM pepstatin A, 762.4 µM leupeptin, 10.6 mM Nα-tosyl-L-arginine methyl ester HCl (Fisher Scientific, T03301G), 10.8 mM tosyl-lysine-chloromethylketone HCl (Fisher Scientific, 50-397-132), 11.3 mM Nα-benzoyl-L-arginine methyl ester carbonate (Fisher Scientific, 50-501-393), 200 µM soybean trypsin inhibitor (Fisher Scientific, NC9065058)), 0.4 mM phenylmethylsulfonyl fluoride (PMSF) and phosphatase inhibitor- PhosStop (Sigma Aldrich, 4906837001). Cell extracts were harvested and cleared by centrifugation. Immunoprecipitation (IP) buffer (50mM HEPES, 100 mM NaCl, 5mM EDTA, and 1% CHAPS (Sigma Aldrich, C3023) with protease inhibitor cocktail, PMSF and PhosStop (as above) was added in a 2:1 ratio to the cell lysate. Samples were incubated with primary antibody or rabbit IgG (control) for 3hr at 4°C and then with Protein A/G PLUS-Agarose (Santa Cruz Biotechnology, sc-2003) beads for 35min with head-to-tail rotation either in the absence or presence of 100 µM CCT blocking peptide SAGRRFIVSPVPE (United Biosystems). Samples were then washed three times with IP buffer before adding 6X SDS sample buffer (0.012% bromophenol blue, 30% glycerol, 10% SDS, 350 mM Tris-Cl, 5% β-mercaptoethanol) and heated at 90°C for 2min. Samples were resolved on 4-20% Mini-PROTEAN® TGX™ Precast Protein Gels (Bio-Rad, 4568096) or 12% polyacrylamide gels and transferred to nitrocellulose membranes. Immunoblotting is as below.

### Immunoblotting

Whole cell lysates containing SDS buffer were homogenized with a 27-G syringe and resolved on 4-20% Mini-PROTEAN® TGX™ Precast Protein Gels (Bio-Rad, 4568096) or 6/10/12% home-made polyacrylamide gels and transferred to nitrocellulose membranes (Fisher Scientific, 45004002). Membranes were washed in Tris-buffered saline (TBS) containing 0.1% Tween® 20 (TBS-T) and blocked with TBS-based blocking buffer (LI-COR). Membranes were incubated with primary antibodies, washed again, incubated with species-specific secondary antibodies. For immunoprecipitation blots, Membranes were washed and blocked as above. Membranes were incubated with primary antibodies, washed again, incubated with species-specific, light chain-specific secondary antibodies (Jackson ImmunoResearch Labs, 115-655-174 and 211-622-171). Blots were analyzed using LI-COR imaging.

### In vitro co-immunoprecipitation

HEK293T were grown in DMEM with 10% FBS. Cells were transfected with pCMV 3xFlag-SPAK 50-545 (human) or Myc-AS160 (residues 193-446). Cells were washed and resuspended in 5 ml cold phosphate-buffered saline (PBS) with 200 µM PMSF and 1:1000 protease inhibitor cocktail (PBSii) as above. The protein fragments were purified as follows: 1 ml of 3xFlag-SPAK supernatant was mixed with 30 µl of anti-Flag magnetic agarose beads (Pierce A36797) and incubated at 4°C for 1hr, washed 3X with 1 ml cold PBSii using magnetic rack, followed by addition of 0.5 ml supernatant Myc-AS160 193-446. The sample was divided in half and 100 µM NH_3_^+^-SAGRRFIVSPVPE-COO^-^ blocking peptide diluted in 25 mM Tris-HCl pH 7.75, 125 mM NaCl was added to one of two and incubated at 4°C for 1hr. Samples were washed 3X with 1 ml cold PBSii. Myc-AS160 193-446 was eluted by addition of 45 µl of 500 µM blocking peptide and 15 µl 5X sample buffer was added to the eluant. Bait and prey proteins were loaded on separate gels (Bio-Rad 4-20% precast gels cat. #4568094). Proteins were transferred to nitrocellulose membrane and immunoblotted as described above. The mouse monoclonal Myc antibody was from clone 9E10. Blots were washed 3 x 20 ml with TBS-T then incubated with 1:5000 IRDye 680 goat anti-Mouse secondary antibody (LI-COR # 926-68070) and analyzed by LI-COR imaging.

### Fluorescence Polarization

3.0 µM His_6_-OSR1 CCT was mixed with 25 nM NH_3_^+^-NLVGRF[DAP-FAM]VSPVPE-COO^-^ (DAP-FAM: 2,3-diaminopropionic acid, unnatural amino acid, conjugated to FAM) in 25 mM Tris-HCl pH 7.75 (at 25° C), 125 mM NaCl, and 1 mM DTT. Unlabeled competing peptides were then added and subjected to a 2-fold stepwise dilution. Fluorescence polarization measurements were carried out on a BioTek Synergy H1 multi-mode plate reader equipped with a fluorescein polarizing filter cube (Em: 485 nm, Ex: 528 nm, 510 nm dichroic mirror). Data was fit to a model of one-site competitive binding to determine K_i_ using GraphPad Prism software.

### Immunofluorescence

SH-SY5Y cells were fixed on glass coverslips (Fisher Scientific, 12-545-80) with 4% paraformaldehyde for 20min at room temperature, washed with PBS and blocked in 10% normal goat serum (Life Technologies, 50-062Z) before incubating with primary antibodies for 1hr at room temperature. After washing with PBS, cells were incubated with an Alexa Fluor® 488 conjugated goat-anti-mouse secondary antibody (Thermo Fisher Scientific, A11029) and Alexa Fluor® 594 conjugated goat-anti-rabbit secondary antibody (Thermo Fisher Scientific, A11037). Slides were mounted with DAPI Fluoromount-G (Thermo Fisher Scientific, 00-4959-52). Immunofluorescent images were acquired on a Zeiss LSM880 inverted confocal microscope (Carl Zeiss, Oberkochen, Germany). Images were deconvolved using AutoQuant® software (Media Cybernetics, USA). The colocalizing pixels were identified and Pearson’s correlation coefficient was determined using Imaris software (Oxford Instruments).

### *In vitro* glucose uptake assay

Cells were placed in DMEM without serum and with DMSO or WNK463 for 2hr and then treated with insulin with or without WNK463 for 20min. A mixture of 0.625 µCi [^3^H]-2-deoxyglucose and unlabeled 2-deoxyglucose (1 mM) was added for 30min at 37°C. After rinsing, cells were lysed, and radioactivity was measured using a scintillation counter. The amount of radioactivity is directly proportional to the rate of glucose utilization [20].

### Surface biotinylation in SH-SY5Y cells

Biotinylation experiments were described previously [179]. Differentiated SH-SY5Y cells in serum-free medium were pre-treated with WNK463 or DMSO for 2hr and treated with insulin with or without WNK463 for 30min. Cells were biotinylated at 4°C with NHS-SS-biotin (0.9 mg/ml) in 10 mM HEPES, 130 mM NaCl, 2 mM MgSO_4_, 1 mM CaCl_2_, 5.5 mM glucose for 15min. After rinsing in 25 mM glycine, cells were lysed in 150 mM NaCl, 50 mM HEPES (pH 7.5), 5 mM EDTA, 1% Triton X-100, 0.2% SDS, and protease inhibitors as above. Lysates were incubated with streptavidin-agarose beads (Pierce Biotechnology) at 4°C overnight. Beads were washed, and biotinylated proteins were extracted by boiling in 60 μl SDS sample buffer with 100 mM dithiothreitol and 5% β-mercaptoethanol. Proteins extracted from the beads (surface) along with total protein were resolved by SDS-PAGE (6% gels). GLUT4 and GAPDH were detected by Western blotting.

### Hippocampal slice culture

Organotypic hippocampal slice cultures were prepared from postnatal day 5 (P5) C57BL/6 mouse strain (Jackson Laboratories, Bar Harbor, ME) and incubated in Minimum Essential Medium (MEM: Gibco, 51200-038) with 5% HyClone Donor Equine Serum (Cytiva, SH30074.03) using previously published protocols [176]. Slices were incubated with WNK463 or DMSO for 1hr. A mixture of 0.625 µCi [^3^H]-2-deoxyglucose and unlabeled 2-deoxyglucose (1 mM) was added for 1min at 37°C. Slices were placed on ice and washed 4X with ice-cold MEM medium. Slices were homogenized in mortar and pestle and radioactivity was measured using a liquid scintillation counter.

### Animal studies

All animal studies were performed according to UTSW Institutional Animal Care and Use Committee guidelines. For studies outlined here, young adult (3-6 months old) male C57Bl/6J mice were used. For harvesting hippocampus from P0-P1 WT C57BL/6J mice for primary cultures, mice were anesthetized by hypothermia followed by decapitation. For other terminal experiments, the method of euthanasia for harvesting tissues was ketamine/xylazine (IP) followed by decapitation for Western blots or [^3^H]-2-deoxyglucose assay. For behavioral and *in vivo* glucose uptake assay, mice were weighed (∼25 g) and orally gavaged daily with 200 µl WNK463 dissolved in 1% DMSO or formulated as a suspension in 0.5:0.5:99 (w:w:w) 2-hydroxypropyl β-cyclodextrin: Pluronic F68: purified water at 6 mg/kg, as indicated.

### *In vivo* glucose uptake assay

1 µCi/g body weight of [^3^H]-2-deoxyglucose was administered into mice by intraperitoneal (IP) injection. After 45min, mice were anesthetized with xylazine/ketamine and decapitated. Brains were removed and immediately placed on ice and hippocampi were dissected and homogenized. The radioactivity was measured using liquid scintillation counter. Plasma samples were taken immediately before mice are sacrificed and radioactivity was analyzed to ensure no significant differences in [^3^H]-2-deoxyglucose [20]. The amount of radioactivity (nCi/g) is directly proportional to the rate of glucose utilization.

### Crude Synaptosome preparation

Crude synaptosomes were prepared as in [180]. C57BL/6J mice were anesthetized as above, brains were removed, and cerebrum dissected and homogenized with mortar and pestle in 0.32 M sucrose, 10 mM HEPES pH 7.4. The homogenates were centrifuged at 1000 x g for 10min at 4 °C and the pellet was removed. The supernatant was centrifuged at 17,000 x g for 30min at 4 °C. This second pellet contained the crude synaptosome fraction and was resuspended in Krebs-Ringer Bicarbonate HEPES buffer (KRBH) buffer containing: 5 mM KCl, 120 mM NaCl, 15 mM HEPES pH-7.4, 24 mM NaHCO_3_, 1 mM MgCl_2_, 2 mM CaCl_2_ (no glucose) and incubated with WNK463 or DMSO for 1hr. A mixture of 0.625 µCi [^3^H]-2-deoxyglucose and unlabeled 2-deoxyglucose (1 mM) was added for 1min at 37°C. Cells were placed on ice and washed 4X with ice-cold KRBH. After sedimentation at 21,000 x g for 15min, radioactivity in the pellet fraction was measured using a liquid scintillation counter.

### Surface biotinylation in brain slices

Biotinylation experiments were performed as described previously [177]. From two mice, 4-5 slices were pooled together randomly for each condition. After a 2-3 h recovery period in aCSF, slices were treated with aCSF containing either DMSO or WNK463 (1 µM) for 45min at 30°C. Slices were treated for 45min with aCSF containing DMSO or WNK463 with or without insulin (10 nM). At the end of the treatment, slices were placed on ice to stop endocytosis and were washed with ice-cold aCSF containing 0.9 mg/ml sulfo-NHS-SS-biotin (Thermo Fisher scientific) for 15min. To quench the biotin reaction, slices were washed once with ice-cold aCSF followed by aCSF containing glycine (25 mM) for 15min and then again with aCSF alone. The hippocampus was dissected from each cerebral slice and homogenized in a modified radioimmunoprecipitation assay (RIPA) buffer containing: 50 mM Tris-HCl, pH 7.4, 1% Triton X-100, 0.1% SDS, 0.5% Na-deoxycholate, 150 mM NaCl, 2 mM EDTA, 50 mM NaH_2_PO_4_, 50 mM NaF, 10 mM Na_4_P_2_O_7_, 1 mM Na_3_VO_4_, and protease inhibitor cocktail as above, followed by sonication. Homogenates were centrifuged at 14,000x g for 10min at 4°C. Protein concentration was measured using BCA Protein Assay (Fisher Scientific, PI23227). 20 µg of protein was removed for total protein measurements. 200 µg protein was then mixed with 200 µl of streptavidin-agarose beads (Thermo Fisher Scientific) by rotating for 2 h at 4°C. The beads were washed twice with 4X volumes of RIPA buffer. Both total and biotinylated (surface) proteins were resolved by 4-20% SDS-PAGE, transferred to nitrocellulose membranes. GAPDH and GLUT4 were detected by Western Blot.

### Insulin Assay

Serum from mice overnight was collected and insulin levels were measured using HTRF Mouse Insulin Serum Detection kit (Revvity).

### Open Field test

Mice were placed in the periphery of a novel open field environment (44 cm x 44 cm, walls 30 cm high) in a dimly lit room (approximately 67 lux) and allowed to explore for 10min. The animals were monitored from above by a video camera connected to a computer running video tracking software (Ethovision XT V-17, Noldus, Leesburg, Virginia) to determine the time, distance moved and number of entries into two areas: the periphery (5 cm from the walls) and the center (14 cm x 14cm). The drug (6 mg/kg) or vehicle was given every day for 3 days prior to testing and was last given 15hr prior to the test and continued until completion of the test. In mice pre-exposed to electric shocks, they received three 0.5 mA foot-shocks at 1min intervals in the fear-conditioning chamber one day before the Open Field test.

### Elevated Plus Maze test

Mice were placed in the center of a black plastic elevated plus maze (each arm 30 cm long and 5 cm wide with two opposite arms closed by 25 cm high walls) elevated 31 cm in a dimly lit room (approximately ∼67 Lux) and allowed to explore for 5min. The animals were monitored from above by a video camera connected to a computer running video tracking software (Ethovision XT V-17, Noldus, Leesburg, Virginia) to determine time spent in the open and closed arms, time spent in the middle, and the number of entries into the open and closed arm. The drug (6 mg/kg) or vehicle was given every day for 3 days prior to testing and was last given 15hr prior to the test and continued until completion of the test. In mice pre-exposed to electric shocks, they received three 0.5 mA foot-shock, 1min interval in the fear-conditioning chamber one day before the Elevated Plus Maze test.

### Fear Conditioning test

Fear conditioning was measured in boxes equipped with a metal grid floor connected to a scrambled shock generator (Med Associates Inc., St. Albans). For training, mice were individually placed in the chamber. After 2min, the mice received 3 tone-shock pairings (30s white noise, 80 dB tone co-terminated with a 2s, 0.5 mA foot-shock, 1min intertrial interval). The following day, memory of the context was measured by placing the mice into the same chambers and freezing was measured automatically by the Med Associates software for 5min. 48hr after training, memory for the white noise cue was measured by placing the mice in a box with altered floors and walls, different lighting, and a vanilla smell. Freezing was measured for 3min, then the noise cue was turned on for an additional 3min and freezing was measured. Mice received oral WNK463 (6 mg/kg) or vehicle 3 days prior to the start of the test and continued until completion of the test.

### Novel Object test

The mice were individually habituated to the test arena (similar to that as the open field test arena) for 2 consecutive days with two identical objects. On the third day, the mice were allowed to explore a different set of similar objects for up to 15min (training). Training concluded when the mouse explored the objects for a total of 30s (both objects combined). Mice that failed to explore the objects for at least 30s were excluded from the study. 6hr after training, a test was performed where one of the familiar objects from testing was replaced by a novel object. Mice received 3 days of daily oral WNK463 (6 mg/kg) or vehicle by oral gavage prior to the start of the test and continued until completion of the test. Time spent exploring the familiar (a) and the novel object (b) was measured. The discrimination index was calculated by the formula (b – a)/(b + a). Mice were excluded from analysis when the total exploration time (b + a) during testing was less than 30s.

### Locomotor Activity test

Mice were placed individually into a clean, plastic mouse cage (18 cm x 28 cm) with minimal bedding. Each cage was placed into a dark Plexiglas box. Movement was monitored by photobeams (Photobeam Activity System, San Diego Instruments, San Diego, CA) for 5hr, with the number of beam breaks recorded every 30min. Mice received 3 days of daily oral WNK463 (6 mg/kg) or vehicle prior to the start of the test.

### Materials, Drugs and Reagents

WNK463 (Selleck Chemicals, S8358), WNK1 siRNA (silencer select custom #1: 5’-3’ CAGACAGUGCAGUAUUCACtt (Thermo Fisher Scientific), anti-Vinculin antibody (Sigma Aldrich, V9131), anti-pOSR1/pSPAK antibody (EMD Millipore, 07-2273), anti-OSR1 polyclonal antibody (Cell Signaling, 3729S), anti-OSR1 monoclonal antibody (VWR, 10624-616), anti-WNK1 antibody (Cell Signaling, 4979S), anti-pAKT S473 antibody (Cell Signaling Technology, 4060S), anti-pAKT T308 antibody (Cell Signaling Technology, 4056S), anti-AKT1 antibody (Cell Signaling Technology, 2920S), anti- GAPDH antibody (Cell signaling Technology, 97166L), anti-GLUT4 polyclonal antibody (ab33780, Abcam), anti-GLUT4 monoclonal antibody (MA5-17176), anti-AS160 antibody (Cell Signaling Technology, 2670S), anti-pAS160 S588 antibody (50-191-485, Fisher Scientific), anti-sortilin antibody (MABN1792, Sigma-Aldrich), anti-Flag antibody (Sigma-Aldrich, F1804). The Myc antibody was a monoclonal from mouse 9E10 (antibody no longer commercially available). Q256 WNK1 antibody was homemade as in [73], Optimem (Invitrogen, 51985-034), Lipofectamine 2000 (Life Technologies, 11668019).

### Statistics and Reproducibility

The data are presented as mean±SEM from at least two-three independent experiments. Micrographs are representative images from at least three experiments. For the quantification of immunofluorescence images, the number of cells used for each representative experiment is indicated and p values between two groups were determined using unpaired t-tests. Single intergroup comparisons between 2 groups were performed with 2-tailed Student’s *t*-test as specifically mentioned in each case. p < 0.05 was considered statistically significant.

### Inclusion and Exclusion criteria

Mice showing at least 50% decrease in phospho-OSR1 levels in the hippocampal lysates were included and used for analysis. No outliers were excluded from the study. Randomization: For inhibitor treatment, mice were randomized before grouping. Blinding: In cases where manual scoring of the behavior was involved, the analysis was performed by an associate blinded to the identity of the mice. Power analysis: On the basis of previous experience of the rodent behavioral core with C57BL/6J mice, animal cohorts of 6-11 mice per group are sufficient to detect differences between groups with a 90% power and a 5% type I error rate.

## Data Sharing

We will follow all NIH policies with respect to sharing reagents, materials, and information with other investigators. Detailed protocols are provided to everyone who requests them. Upon publication, this manuscript will be submitted to the National Library of Medicine’s PubMed Central as outlined by NIH policy.

## Acknowledgements

The authors thank the members of Cobb and Huber labs for valuable suggestions, and Dionne Ware for administrative assistance. We would like to thank Dr. Joseph Albanesi, Department of Pharmacology, University of Texas Southwestern Medical Center, for discussions and suggestions related to the manuscript and figures. We would like to thank Steve Stippec (Cobb lab) for his help with molecular cloning of constructs used in this study. We would like to thank Gemma Molinaro and Julia Wilkerson (Huber lab) for providing technical support with primary cortical neurons/glial cell culture, hippocampal slice culture and preparation of hippocampal slices for surface biotinylation experiments. We would also like to thank Barbara Barylko, Department of Pharmacology, University of Texas Southwestern Medical Center for assistance with crude synaptosome preparation and Jenna Jewell, Department of Molecular Biology, University of Texas Southwestern Medical Center, for HEK293 cells. We also would like to thank Julia Watkins at UNT Heath, Fort Worth for her technical assistance in revising the manuscript. These studies were supported by NIH K99AG075161-01A1 to ABJ, NIH R01 HL147661 to MHC, Welch Foundation grant I1243 to MHC, and 1R37NS114516-01A1 to KH. Behavioral experiments were performed in collaboration with Rodent Behavior Core, (supported in part by the Peter O’Donnell Jr. Brain Institute, University of Texas Southwestern Medical Center). The authors would also like to acknowledge the assistance of the UT Southwestern Live Cell Imaging Facility, a shared resource of the Harold C. Simmons Comprehensive Cancer Center, supported in part by an NCI Cancer Center Support Grant, 1P30 CA142543-01 and NIH Shared Instrumentation Award 1S10 OD021684-01 to Dr. Kate Luby-Phelps (LSM880 Airyscan).

## Author Contributions

**ABJ:** Conceptualized, supervised, designed and performed experiments, performed analysis, wrote manuscript, generated initial figures; **AA**: performed experiments; **DB**: performed experiments; **CAT**: performed experiments; **SGB**: Experiment design; **KH**: Supervision and acquired funding; **MHC**: Supervision, acquired funding, edited manuscript.

## Supplementary Figure Legends

**Supplementary Figure 1.**
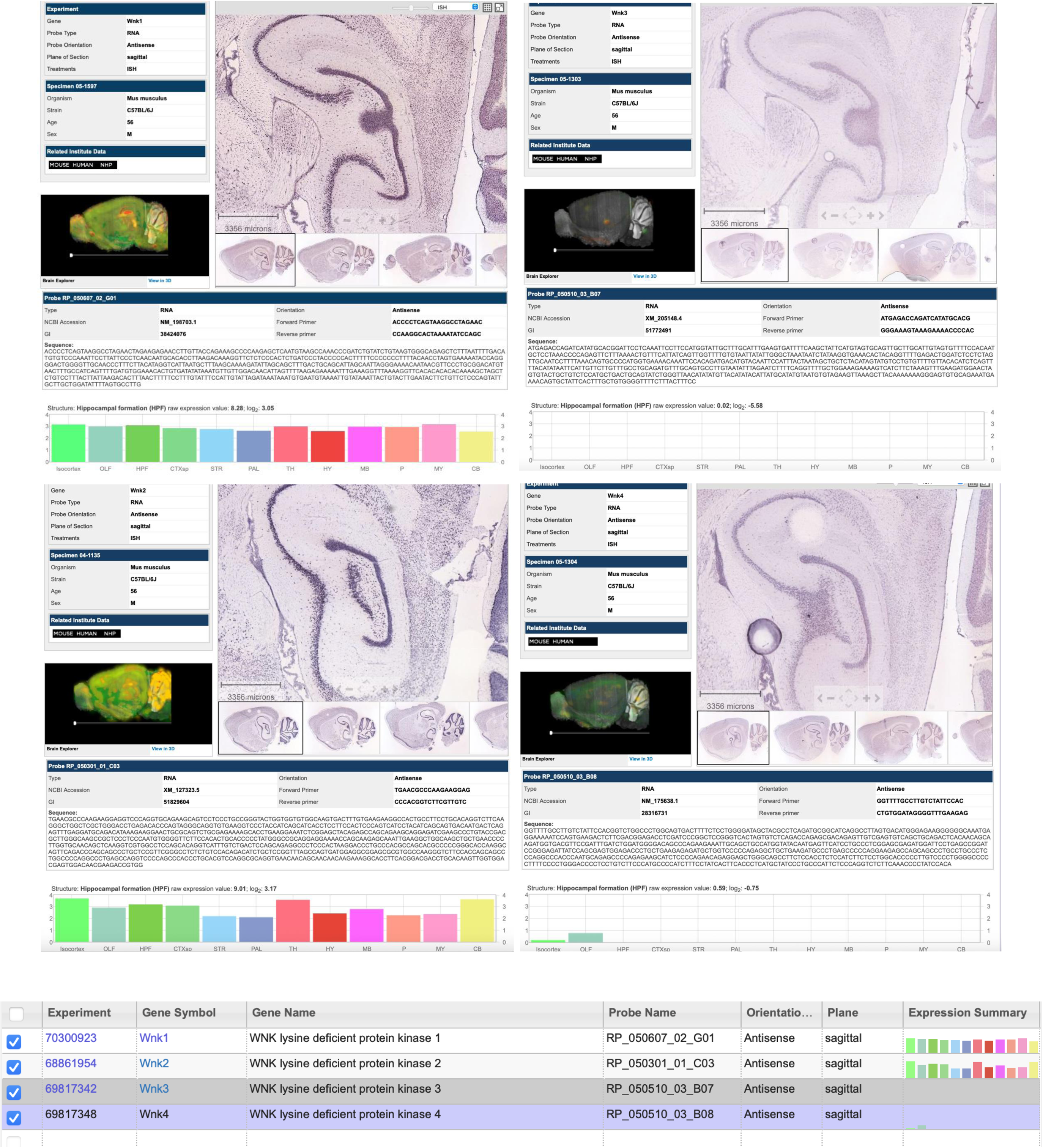
**A)** Mouse in situ hybridization data curated from Allen Brain Atlas shows WNK 1-4 expression in the hippocampus (**Figure S1A**).

**Supplementary Figure 2.**
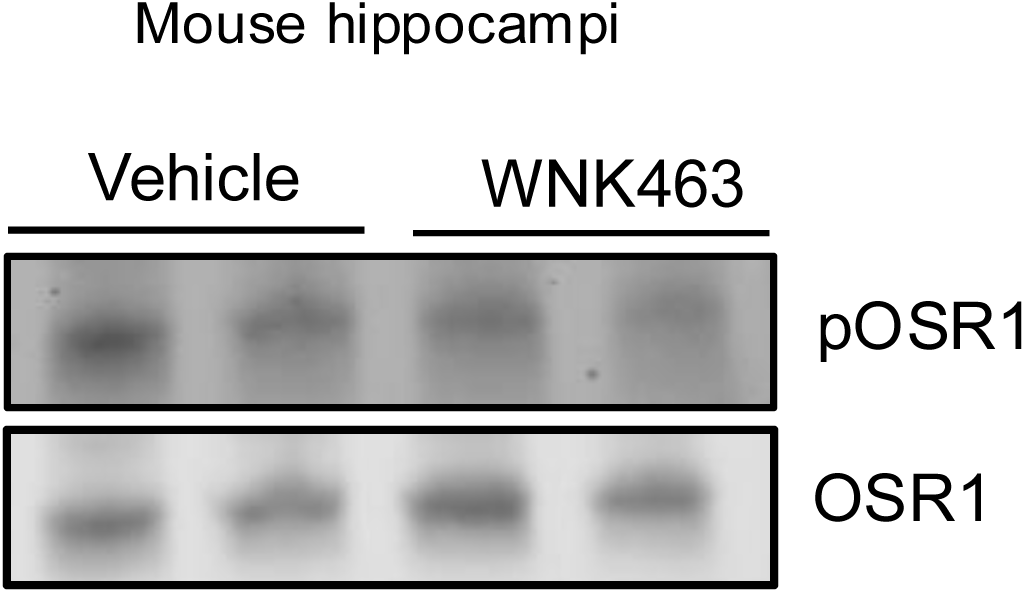
**A)** Representative Western blots (for quantification shown in Figure 1B) shows OSR1 levels in excised hippocampal tissue from mice treated with vehicle or WNK463 6 mg/kg for 3 days.

**Supplementary Figure 3.**
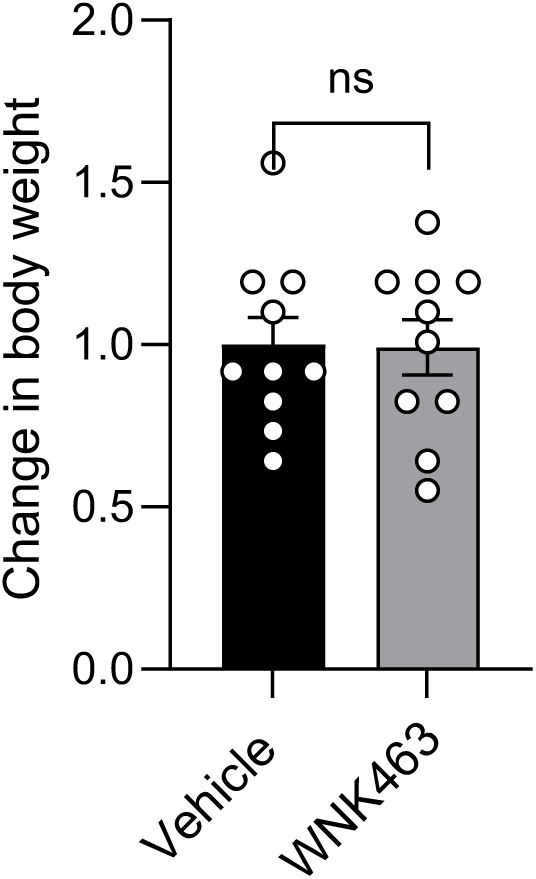
**A)** Graph showing change in body weights of C57BL/6J mice treated with vehicle or WNK463 (PO; 6mg/kg) for 1 week; n=10. Data are represented as Mean±SE; analyzed by unpaired two-tailed Student’s *t*-test. ns: non-significant; p>0.05.

**Supplementary Figure 4.**
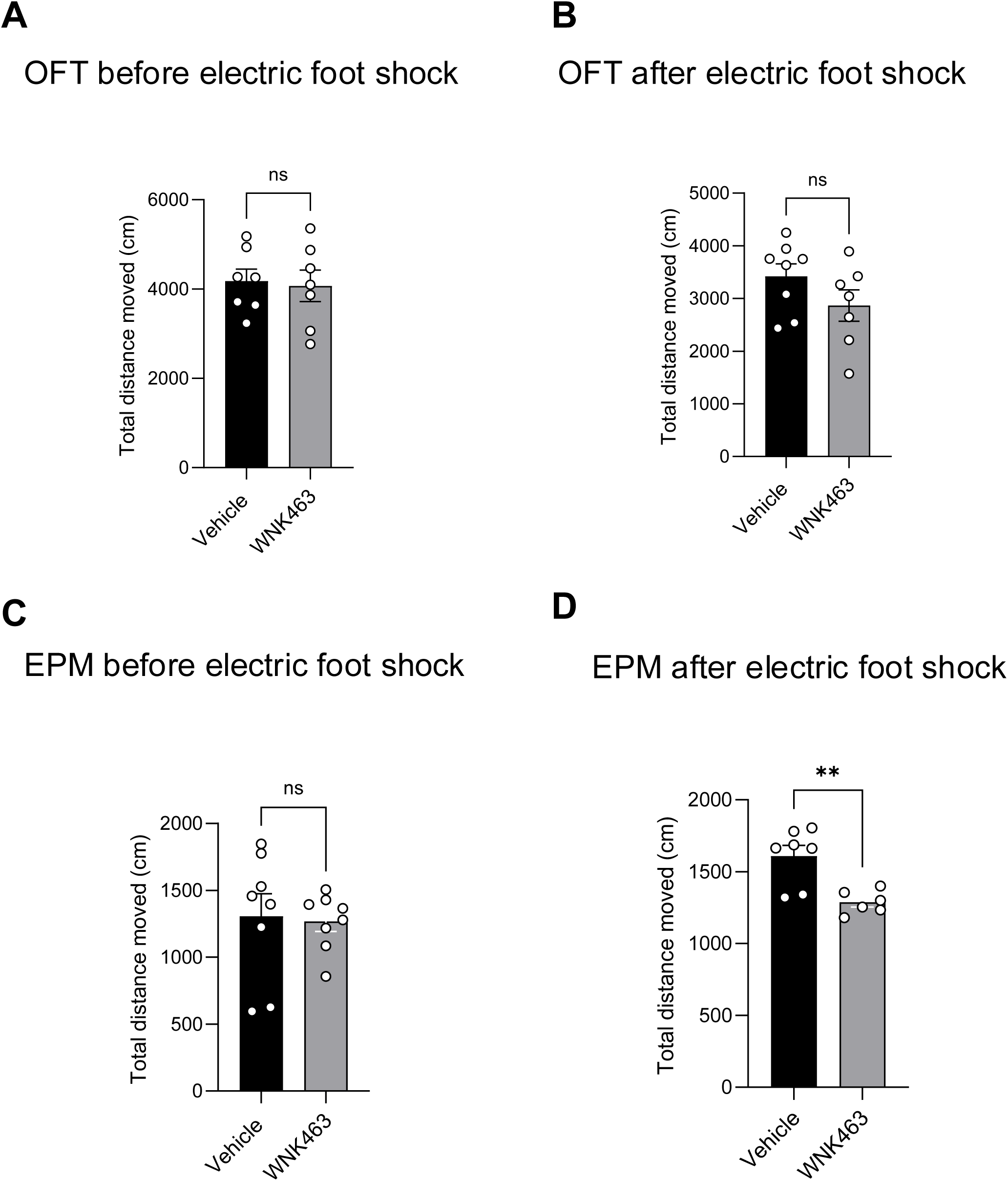
**A)** Quantification of total distance moved (cm) in the Open Field test in mice administered with WNK463 (PO; 6 mg/kg) or vehicle; n=7. **B)** Quantification of total distance moved (cm) in the Open Field test in mice administered with WNK463 (PO; 6 mg/kg) (n=7) or vehicle (n=8) in mice pre-exposed to electric foot-shocks. **C)** Quantification of total distance moved (cm) in the Elevated Plus Maze test in mice administered with WNK463 (PO; 6 mg/kg) or vehicle; n=8. **D)** Quantification of total distance moved (cm) in the Elevated Plus Maze test in mice administered with WNK463 (PO; 6 mg/kg) or vehicle in mice pre-exposed to electric foot-shocks; n=7. Data are represented as Mean±SE; analyzed by unpaired two-tailed Student’s *t*-test. *p<0.05, **p<0.005 and *** p<0.0005. ns: non-significant; p>0.05.

**Supplementary Figure 5.**
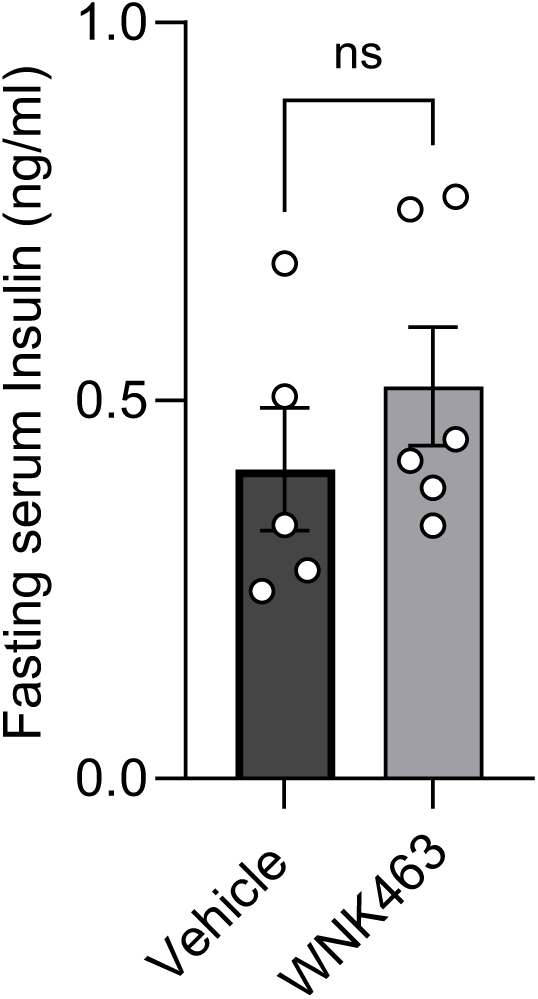
**A)** Quantitative graph shows fasting serum insulin from mice treated with vehicle or oral WNK463 (6mg/kg) for 3 days; n=5. Data are represented as Mean±SE; analyzed by unpaired two-tailed Student’s *t*-test. ns: non-significant; p>0.05.

**Supplementary Figure 6.**
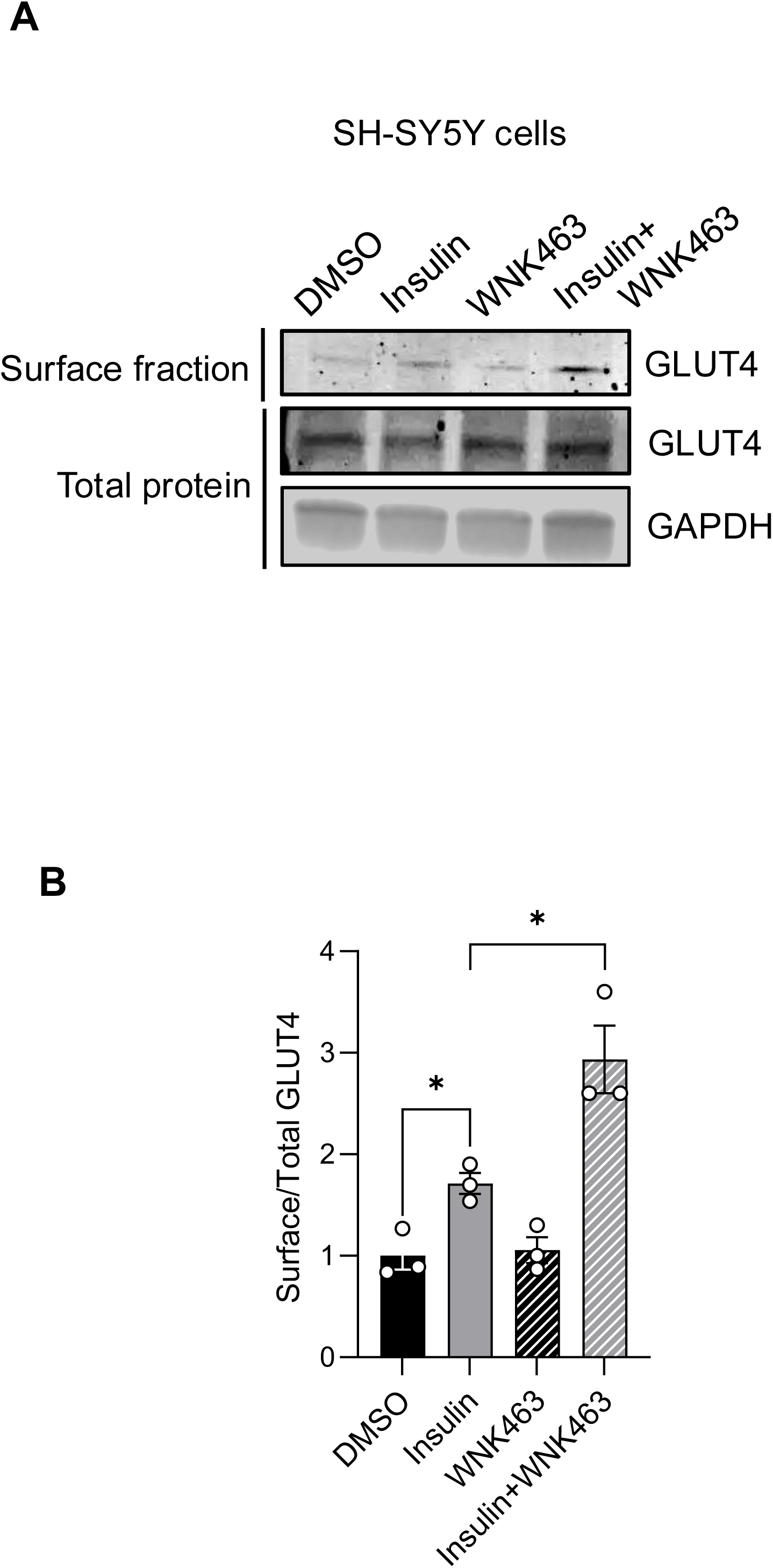
**A)** Representative Western blot shows surface and total GLUT4 protein fraction in differentiated SH-SY5Y cells treated with insulin (10 nM) and/or WNK463 (1 µM). **B)** Corresponding quantification of ‘A’ shows enhanced surface GLUT4 (measured as a fraction of total GLUT4) with WNK463 ± insulin treatment; n=3. Data are represented as Mean±SE; analyzed by unpaired two-tailed Student’s *t*-test. *p<0.05, **p<0.005 and *** p<0.0005.

**Supplementary Figure 7.**
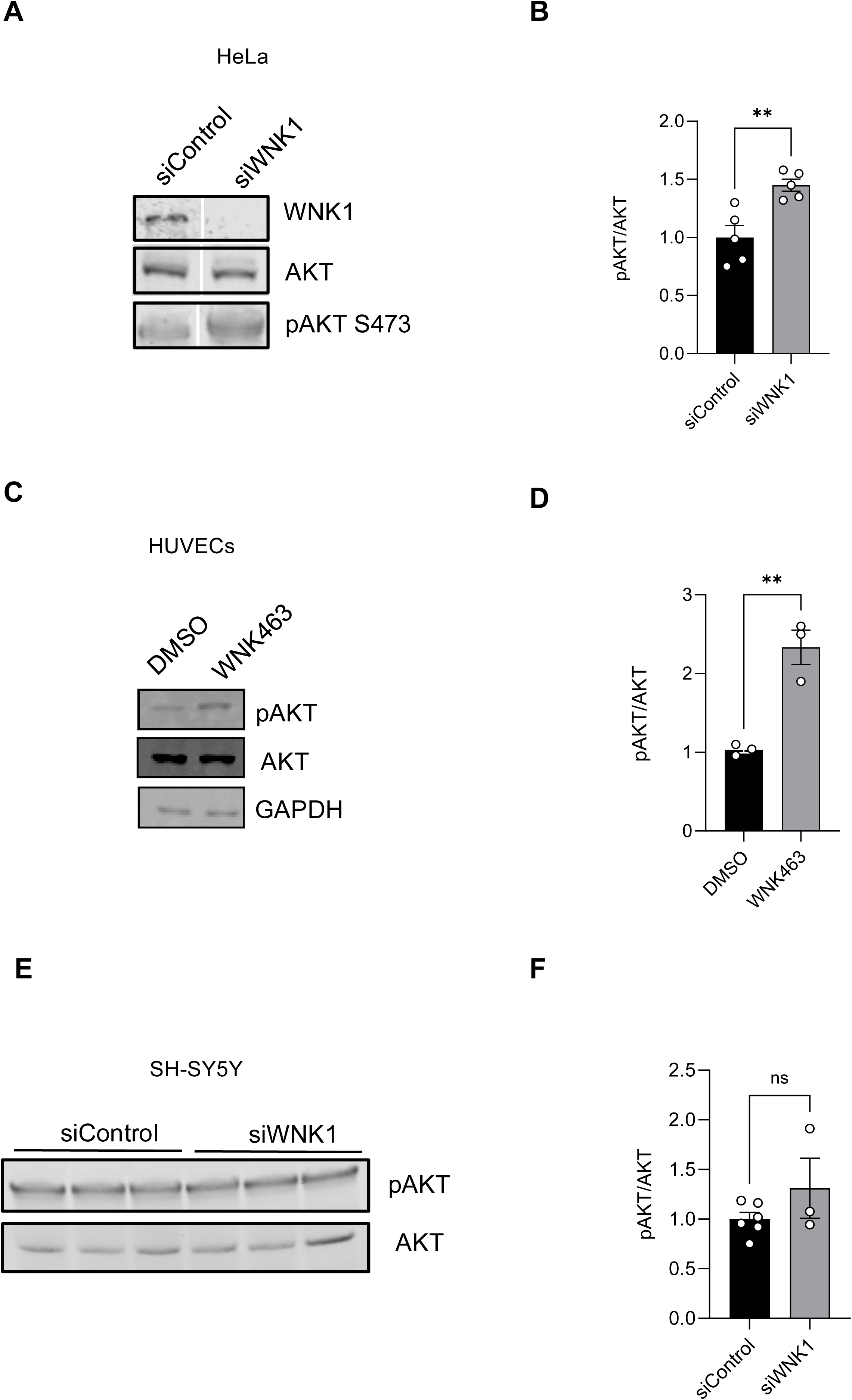
**A)** Representative Western blot shows enhanced pAKT levels upon treatment with WNK1 siRNA in HeLa cells; n=5. **B**) Corresponding quantification of ‘A’. **C**) Representative Western Blot shows elevated pAKT levels in primary Human Umbilical Vein Endothelial cells upon WNK463 treatment; n=3. **D**) Corresponding quantification of ‘C’. **E**) Representative Western blot shows pAKT levels in differentiated SH-SY5Y cells treated with WNK1 siRNA; n=3. **F**) Corresponding quantification of ‘E’. Data are represented as Mean±SE; analyzed by unpaired two-tailed Student’s *t*-test. ns: non-significant; p>0.05, *p<0.05, **p<0.01.

**Supplementary Figure 8.**
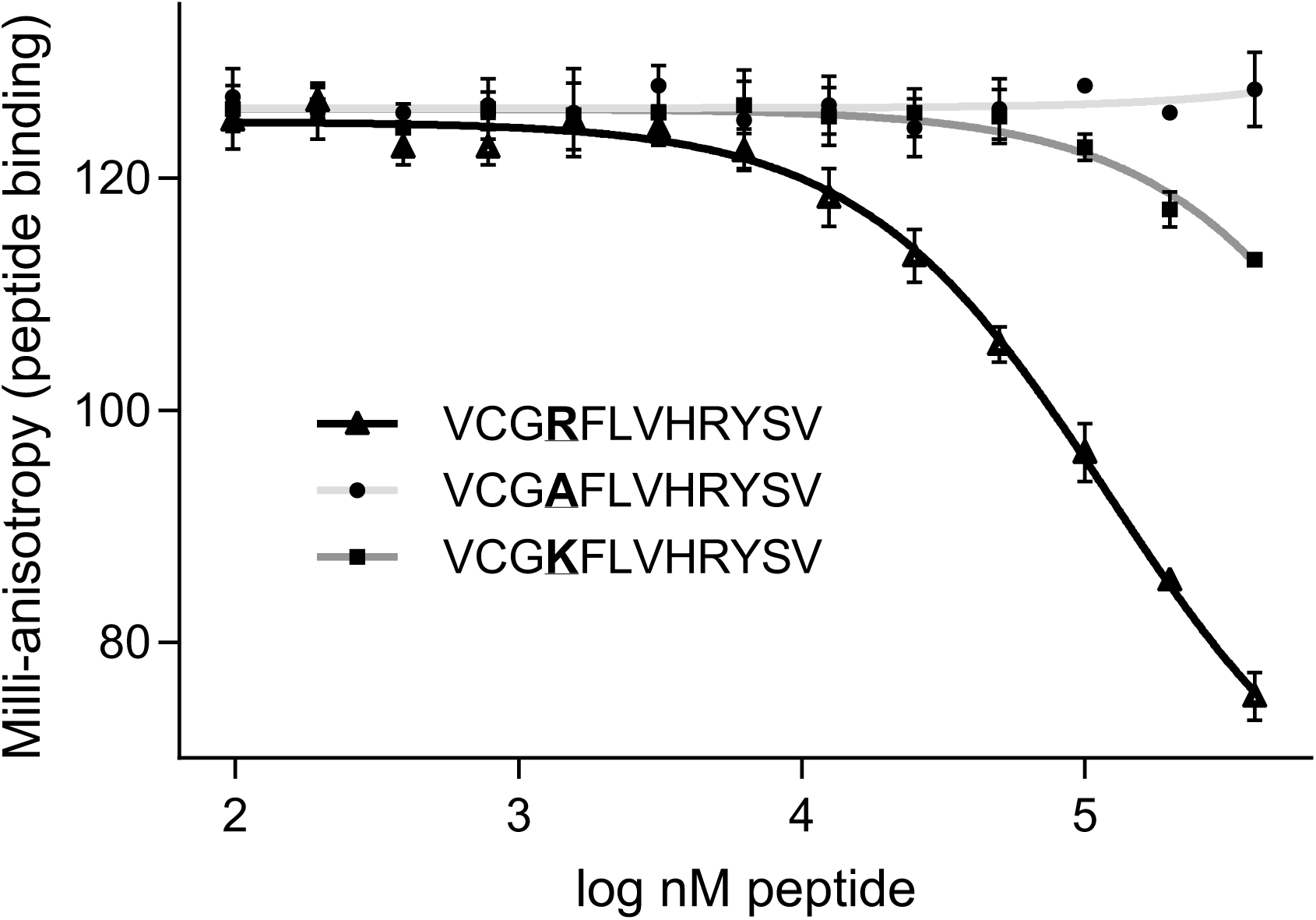
**A)** Graph representing the binding affinities of WT- sortilin and sortilin mutant peptides with OSR1/SPAK CCT determined by fluorescence anisotropy. Unlabeled peptides displace labeled peptide (NH3^+^-NLVGRF-[DAP-FAM]-VSPVPE-COO^−^) [diaminopropionic acid (DAP)]. Labeled peptide held constant at 25 nM, OSR1 CCT held constant at 3.0 μM; n=3.

